# Protein homeostasis imposes a barrier on functional integration of horizontally transferred genes in bacteria

**DOI:** 10.1101/025841

**Authors:** Shimon Bershtein, Adrian W.R. Serohijos, Sanchari Bhattacharyya, Michael Manhart, Jeong-Mo Choi, Wanmeng Mu, Jingwen Zhou, Eugene I. Shakhnovich

## Abstract

Horizontal gene transfer (HGT) plays a central role in bacterial evolution, yet the molecular and cellular constraints on functional integration of the foreign genes are poorly understood. Here we performed inter-species replacement of the chromosomal *folA* gene, encoding an essential metabolic enzyme dihydrofolate reductase (DHFR), with orthologs from 35 other mesophilic bacteria. The orthologous inter-species replacements caused a marked drop (in the range 10-90%) in bacterial growth rate despite the fact that most orthologous DHFRs are as stable as *E.coli* DHFR at 37° C and are more catalytically active than *E. coli* DHFR. Although phylogenetic distance between *E. coli* and orthologous DHFRs as well as their individual molecular properties correlate poorly with growth rates, the *product* of the intracellular DHFR abundance and catalytic activity (*k*_*cat*_/K_M_), correlates strongly with growth rates, indicating that the drop in DHFR abundance constitutes the major fitness barrier to HGT. Serial propagation of the orthologous strains for ∼600 generations dramatically improved growth rates by largely alleviating the fitness barriers. Whole genome sequencing and global proteome quantification revealed that the evolved strains with the largest fitness improvements have accumulated mutations that inactivated the ATP-dependent Lon protease, causing an increase in the intracellular DHFR abundance. In one case DHFR abundance increased further due to mutations accumulated in *folA* promoter, but only after the *lon* inactivating mutations were fixed in the population. Thus, by apparently distinguishing between self and non-self proteins, protein homeostasis imposes an immediate and global barrier to the functional integration of foreign genes by decreasing the intracellular abundance of their products. Once this barrier is alleviated, more fine-tuned evolution occurs to adjust the function/expression of the transferred proteins to the constraints imposed by the intracellular environment of the host organism.

## Author Summary

Horizontal gene transfer (HGT) is central to bacterial evolution. The outcome of an HGT event (fixation in a population, elimination, or separation as a subdominant clone) depends not only on the availability of a new gene but crucially on the fitness cost or benefit of the genomic incorporation of the foreign gene and its expression in recipient bacteria. Here we studied the fitness landscape for inter-species chromosomal replacement of an essential protein, dihydrofolate reductase (DHFR) encoded by the *folA* gene, by its orthologs from other mesophilic bacteria. We purified and biochemically characterized 33 out of 35 orthologous DHFRs and found that most of them are stable and more catalytically active than *E. coli* DHFR. However, the inter-species replacement of DHFR caused significant fitness loss for most transgenic strains due to low abundance of orthologous DHFRs in *E. coli* cytoplasm. Laboratory evolution resulted in an increase in orthologous DHFR abundance leading to a dramatic fitness improvement. Genomic and proteomic analyses of “naive” and evolved strains suggest a new function of protein homeostasis to discriminate between “self” and “non-self” proteins, thus creating fitness barriers to HGT.

## Introduction

Horizontal gene transfer (HGT) is a major force in bacterial evolution [1-3]. Comparative genomic analyses show that HGT events can be broadly classified into three types: a) acquisition of a new gene not present in the taxa, b) acquisition of an orthologous gene in addition to the endogenous chromosomal copy, and c) direct chromosomal replacement of a gene by its ortholog from other species (also known as xenologous horizontal gene transfer) [1]. Koonin *et al.* also found that all three HGT types are approximately equally common and represent an efficient mechanism for rapid evolution and/or adaptation to new niches [1].

The genetic mechanisms responsible for the horizontal transfer of foreign genes (*i.e.*, transformation of naked DNA, conjugation, and viral transduction) are well characterized [4-6]. However, the material transfer of DNA from other bacterial species is only an initial step. The evolutionary fate of an HGT event (fixation, elimination by purifying selection, or persistence as a subdominant clone) depends on the fitness benefit or cost of the newly acquired gene. Previous studies on these fitness effects have arrived at apparently inconsistent conclusions. For example, Sorek *et al.* [7] expressed multiple proteins from 79 prokaryotic genomes in an expression vector under control of an inducible promoter and measured the ensuing fitness effects in *E. coli*. They found that expression of many foreign proteins is detrimental to the *E. coli* host and attributed the fitness cost to a gene dosage-related toxicity. Lind *et al.* found that inter-species chromosomal replacement of three native genes encoding ribosomal proteins in *S. typhimurium* was detrimental to fitness, apparently due to low expression of transferred proteins [8]. Although these studies showed that many HGT events incur fitness costs, they did not provide mechanistic or molecular explanations of why this was the case. Meanwhile, other studies have argued that HGT is predominantly neutral rather than deleterious. Insertion of random DNA fragments from other bacteria in the *Salmonella* chromosome showed no significant fitness effect for ∼90% of the inserts [9]. Introduction of foreign and complex subunits in *E. coli* also showed no loss in fitness [10].

The apparent controversies on the nature of the fitness landscape of HGT events can be attributed to several challenges. A) *Pleiotropy at the molecular level.* The starting genetic material has a broad distribution of molecular and sequence properties that are not entirely independent (e.g., potential effect of GC-content on RNA stability [11] that could affect transcription/translation, and of protein folding stability and activity [12] that could affect function). B) *Pleiotropy at the cellular level.* Beyond the foreign gene’s immediate functional context, HGT may affect or be affected by other cellular factors, such as protein-protein, metabolic, or regulatory interaction networks [13-16]. Another example is protein homeostasis (proteostasis) machinery, which maintains the integrity of the proteome through assisted folding and degradation and is known to buffer against the deleterious effects of mutations [17-19]. However, the actual effect of proteostasis on horizontal gene transfer is not yet known. C) *Time and length scale in evolution*. Similarly to mutations, HGT events can be accompanied by immediate and transient responses of the cell that are particularly hard to detect using comparative genomics, because it analyzes HGT that has survived selection over long evolutionary time scales. Altogether, these challenges need to be addressed to understand the fitness landscape of HGT and the cellular responses that lead to the subsequent accommodation or rejection of a foreign gene.

Here we aimed at a molecular-and systems-level mechanistic description of the origin of the fitness cost associated with HGT immediately upon its occurrence, as well as after a period of experimental evolution. In particular, we sought to develop an experimental system that allows full control over the molecular properties of the transferred gene (**Fig 1A**). Our focus is on the functional barriers to HGT emerging at the protein level rather than genomic barriers affecting transcription and translation. To this end, we used the essential gene *folA* encoding dihydrofolate reductase (DHFR) as a model. DHFR catalyzes an electron-transfer reaction to form tetrahydrofolate, a carrier of single-carbon functional groups utilized in central metabolism, including *de novo* purine biosynthesis, dTTP formation, and methionine and glycine production [20]. DHFR is also an important target of antifolate therapy by trimethoprim (TMP), a competitive inhibitor that binds with high specificity to the active site of bacterial enzymes [21]. Additionally, comparative genomics studies have demonstrated that HGT plays an important role in the evolution of the folate metabolic pathway, including the spread of antifolate resistance [22,23]. Moreover, DHFR is an essential enzyme in *E. coli* with a relatively low basal expression level (approximately 40 copies per cell on average [24]), and its activity is linked to bacterial fitness in a dosage-dependent manner [17]. As such, DHFR is a convenient model to study the HGT-related fitness effects.

**Fig 1.**
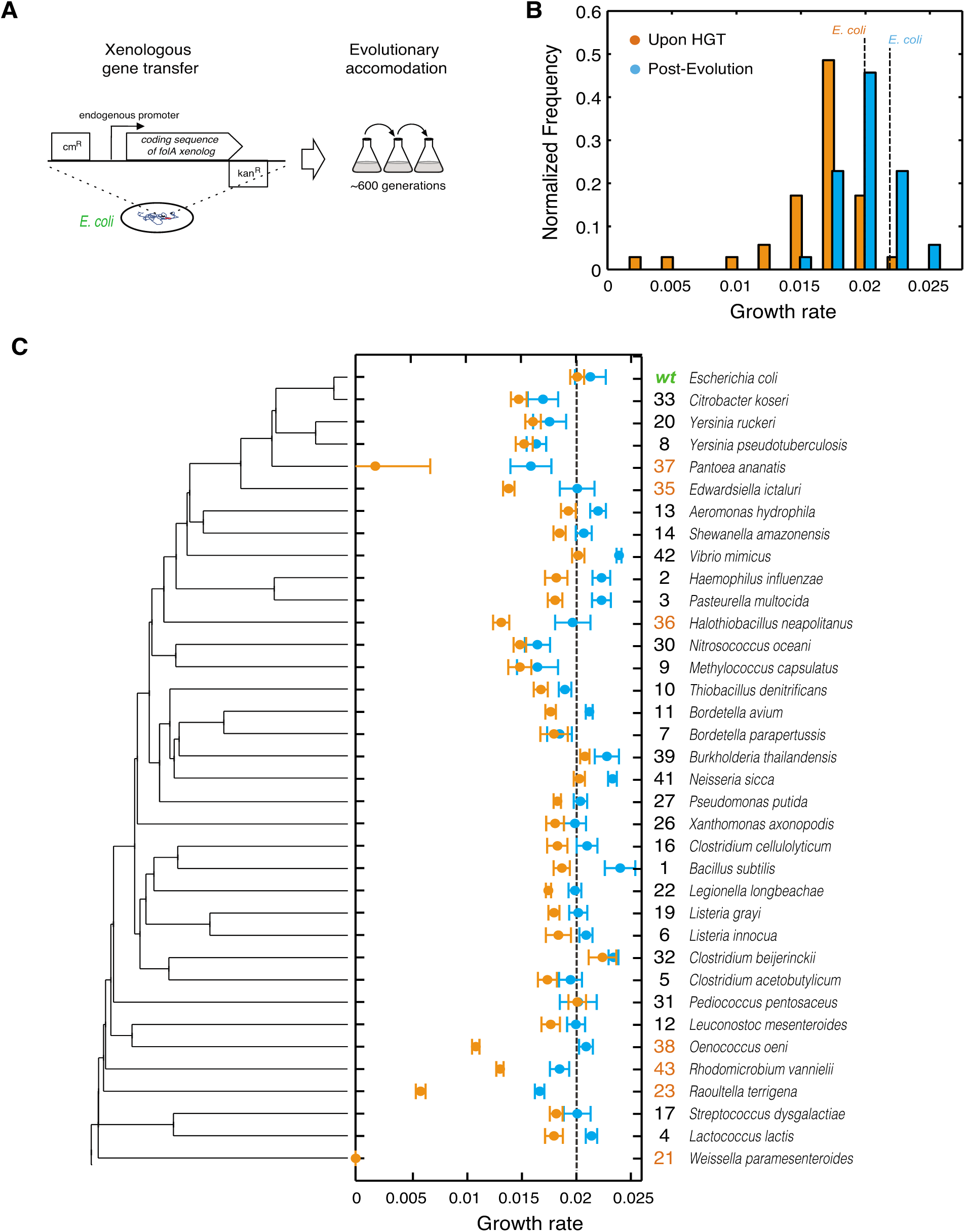
The barrier to horizontal transfer of orthologous DHFR proteins is alleviated by experimental evolution. A) ORF of *folA* gene encoding DHFR in the *E. coli* chromosome is replaced with orthologs from 35 other mesophiles, while preserving the endogenous promoter. The strains carrying the orthologous DHFR replacements are evolved for 31 serial passages (∼600 generations) under standard conditions. B) Distribution of the growth rates before (immediately upon HGT) and after evolution experiment. Growth rate of the WT strain with *E. coli* DHFR is marked with a dashed line (see also **S3 Table**). Kolmogorov-Smirnov (KS) test indicates that pre-and post-evolution populations are significantly different with respect to their growth rates (p-value <10^−10^). While 31 out of the 35 naive strains (88%) have lower growth rates than WT, 30% of the post-evolution strains have higher growth rates than WT. C) Growth rates before and after evolution for each strain as a function of its DHFR’s position in the phylogeny. Color scheme is similar to panel (B). Strains are ordered according to the phylogenetic tree on the left. On the right we show an ID number for each strain (used throughout the text) and the original species carrying that DHFR ortholog. We highlight in orange the ID numbers of strains that experience severe fitness drop (30% and lower) upon DHFR replacement. Error bars represent standard deviation of 4 independent measurements.

We experimentally mimicked multiple HGT events by replacing the *folA* gene on the *E. coli* chromosome with its orthologs from 35 phylogenetically diverse mesophiles. This collection of orthologs explores a broad distribution of protein sequences and biophysical properties. We found that immediately after HGT the fitness effects were largely deleterious, whereas after the experimental evolution fitness has improved markedly. Using biochemical and genetic analysis, whole-genome sequencing, and proteomics, we show that the mechanistic origin of the barriers to HGT is the global response of the protein homeostasis machinery. Altogether, this work provides molecular and cellular level insights into the origin of the barriers that shape the fitness landscapes of HGT events.

## RESULTS

### Orthologous replacement of DHFR is detrimental to bacterial fitness

We initially identified 290 orthologous DHFR sequences from mesophilic bacteria and selected 35 diverse sequences with amino acid identity to *E. coli* DHFR ranging from 29% to 96% (**Fig 1C, S1 Table**, and **Materials and Methods**). First, we sought to minimize the contributions from confounding factors that mostly affect transcription and translation of replaced genes, such as GC content [25], codon-usage pattern [26,27], specific loci at which chromosomal incorporations occur, and the copy number of the transferred genes [6,11]. The amino acid sequences of the chosen 35 orthologous DHFRs were converted into DNA sequences using the codon signature of *E. coli*’s *folA* gene (**Materials and Methods**, **S2 Table**). We used the λ-red recombination system [28] to replace the open reading frame (ORF) of *folA* with the synthetic DNA sequences, while preserving *E*. *coli*’s wild-type *folA* promoter (see **Materials and Methods**). Thus, the resulting 35 strains carrying the orthologous DHFR gene replacements are identical with respect to the chromosomal location of the *folA* gene and the mode of regulation of their DHFR expression. In addition, they have similar GC content and codon usage signature (**S1 and S2 Tables**).

We assayed the fitness of the resulting HGT strains by measuring their growth rates at 37°C (this condition was consistently used throughout the work) (see **Materials and Methods**, **Fig 1B,C** and **S3 Table**). We found that *E. coli* fitness (here and below we use the terms fitness and growth rate interchangeably) is very sensitive to the orthologous replacements of its DHFR. Growth rates are lower than wild-type (WT) *E. coli* in 31 out of 35 strains, with six strains (DHFR-23, 35, 36, 37, 38 and 43; highlighted in **Fig 1C**) exhibiting a severe fitness loss of 70-85%. DHFR-21 (from *W. paramesenteroides*) did not grow at all under the conditions of the experiments and was omitted from further analysis. Surprisingly, we found no significant correlation between growth rates of the HGT strain and the evolutionary distance between DHFR orthologs, measured as % of amino acid identity relative to *E. coli* DHFR (Spearman r=0.16; p-value=0.4) (**S1A Fig**), thus, challenging the notion that sequence similarity between endogenous and transferred genes facilitates horizontal gene transfer.

### Orthologous DHFRs are not toxic to the cell

A potential explanation for the behavior of strains that exhibit most detrimental effect of HGT, namely, DHFR-23, 35, 36, 38, and 43 is that these new proteins are inherently toxic to the cell, as previously found for many horizontally transferred genes [7]. To determine whether expression of orthologous DHFRs is toxic to the cell, we followed the approach of Sorek and co-workers [7] by transforming WT *E. coli* cells with pBAD plasmids expressing the orthologous DHFRs under the arabinose-controlled promoter (**Materials and Methods**). We found that none of the orthologous DHFRs caused a significant drop in fitness (S2A,B Fig). Alternatively, we transformed the severely affected HGT strains (DHFR-23, 35, 36, 37, 38, and 43) with pBAD plasmids expressing WT DHFR (**Materials and Methods**). As shown in **S2C Fig**, complementation with WT DHFR completely restored fitness. Thus, the fitness cost associated with the HGT of DHFR-23, 35, 36, 37, 38, and 43 is not caused by toxicity, but by the loss of DHFR function in the cells.

### Experimental evolution allows bacteria to cross HGT fitness barriers

The high fitness cost of the orthologous replacements of *E. coli* DHFR demonstrates the existence of a molecular constraint (“a barrier”) to HGT. To determine whether the evolutionary process can traverse this barrier, we conducted high-throughput serial passaging of the HGT strains (see **Materials and Methods**). Overall, we performed 31 passages which amount to ∼600 generations for the WT strain. Growth rate measurements after the evolution experiment show that orthologous strains have substantially improved their fitness (**Fig 1C** and **S2 Table**). Moreover, ∼30% of the strains grew as well as or better than WT after the evolution experiment (**Fig 1B,C**). The improvement in growth rates was especially dramatic amongst strains that experienced the most severe fitness loss upon HGT (DHFR-23, 35, 36, 37, 38 and 43; highlighted in **Fig 1C**). Thus, the molecular constraints to horizontal transfer of the DHFR coding genes were largely alleviated during experimental evolution.

### Distribution of the molecular and cellular properties of orthologous DHFRs

To determine whether the mechanisms underlying the barrier to HGT are linked to the molecular properties of the horizontally transferred proteins, we looked at the important *in vitro* biophysical properties of the orthologous DHFRs (**Fig 2A,B**) (see **Materials and Methods**). The thermodynamic stabilities of orthologous DHFRs quantified by *T*_*m*_ (mid-transition temperature of thermal unfolding) fall within the range expected for mesophilic proteins (42-63°C) (**Fig 2A** and **S1 Table**) [29]. Approximately 20% of DHFRs are more thermodynamically stable than *E. coli* DHFR (**Fig 2A**). **Fig 2B** shows that the catalytic activity (*k*_*cat*_/*K*_*m*_) of orthologous DHFRs varies widely, with approximately ∼70% of enzymes having activities that are comparable to or better than *E. coli* DHFR. Some enzymes, in fact, exhibit almost 100-fold higher activity than *E. coli* DHFR (**Fig 2B, S1 Table**). No significant correlation was found between the *in vitro* properties and fitness upon the HGT (**S3A,C Fig**).

**Fig 2.**
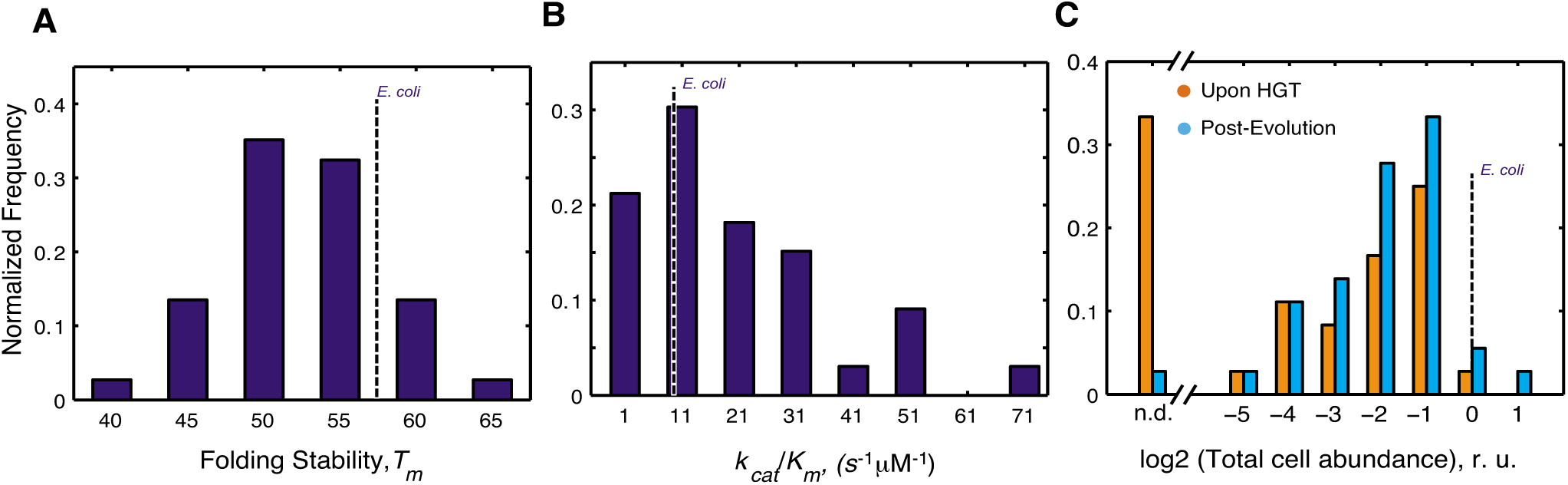
Distribution of molecular and cellular properties of orthologous DHFRs. A) Distribution of T_m_ values of the purified DHFR proteins assessed by thermal unfolding in a Differential Scanning Calorimeter (see **Materials and Methods**). The proteins span wide range of stability between 42-63°C, as expected for mesophilic proteins, with ∼80% of proteins having higher stability than *E.coli* DHFR. (see also **S1 Table**). B) Distribution of catalytic activity (*k*_*cat*_/K_M_) of the purified DHFRs (see **Materials and Methods**, and **S1 Table**). ∼70% purified DHFR proteins were found to have activities that are comparable to or better than E. coli DHFR. C) Intracellular abundance (measured in total cell lysate) of DHFR before and after evolution experiment assessed by Western Blot with polyclonal anti-His antibodies (see **Materials and Methods**). Abundance is expressed relative to that of *E. coli* DHFR. Strains for which abundances were too low to be detected are denoted as n.d. After the evolution experiment, a large number of strains have detectable abundance (∼97% compared to ∼70% before evolution) (**S1 Table** and S5 Fig) Overall, there is a significant shift in the abundance distribution for all proteins after evolution experiment (KS p-value=0.049).

Effective metabolic turnover, however, is not only a function of enzymatic activity, but also of intracellular abundance. Thus, we next measured the total intracellular abundances of DHFR in all strains (see **Materials and Methods**). We found that immediately upon HGT (before the evolution experiment), soluble and total abundances of orthologous DHFRs in all strains were lower than WT DHFR, and for ∼35% of the strains, DHFR abundance was bordering the detection limit (**Fig 2C, S3 Table, S5 Fig**). After the evolution experiment, however, we observed a significant increase in total DHFR abundances (**Fig 2C, S3 Table, S5 Fig**) (Kolmogorov-Smirnov (KS)-test, p-value=0.049). We found that strains with undetectable DHFR immediately upon HGT (DHFR-23, 35, 36, 37, 38 and 43) increased their abundances to detectable levels after the evolution experiment (**Fig 2C, S3 Table, and S5 Fig**). This finding suggests that the low intracellular abundance of the foreign DHFR proteins is the direct manifestation of the HGT barrier. Moreover, alleviation of the barrier during experimental evolution leads to an increase in functional copies of DHFR.

### Dosage-dependent fitness landscape of HGT

To understand how HGT shapes the fitness landscape of *E. coli* cells, we projected the growth rates of the HGT strains onto the catalytic activity of the DHFR proteins (**Fig 3A,B**). No correlation was observed immediately upon HGT, primarily because several of the strains (DHFR-23, 35, 36, 38, and 43) exhibited severe fitness drops despite carrying orthologous DHFR proteins with *k*_*ca*t/_*K*_*m*_ values that are comparable to or higher than *E. coli* DHFR (**Fig 3A**). We were unable to purify the severely deleterious DHFR-37, thus there is no *k*_*cat*_/*K*_*m*_ value for this protein. After the evolution experiment, however, growth of the severely affected HGT strains improved dramatically (by ∼60-90%). This change resulted in a statistically significant correlation between catalytic activity and fitness among *all* the points (Spearman r= 0.57, p-value = 0.0007; **Fig 3B, 4A**). We note that the theoretical fit in **Fig 3** (green lines) excludes DHFR-23, 35, 36, 38, and 43; nonetheless, after the evolution experiment, these initial outliers cluster around the theoretical prediction (**Fig 3B**, orange points).

**Fig 3.**
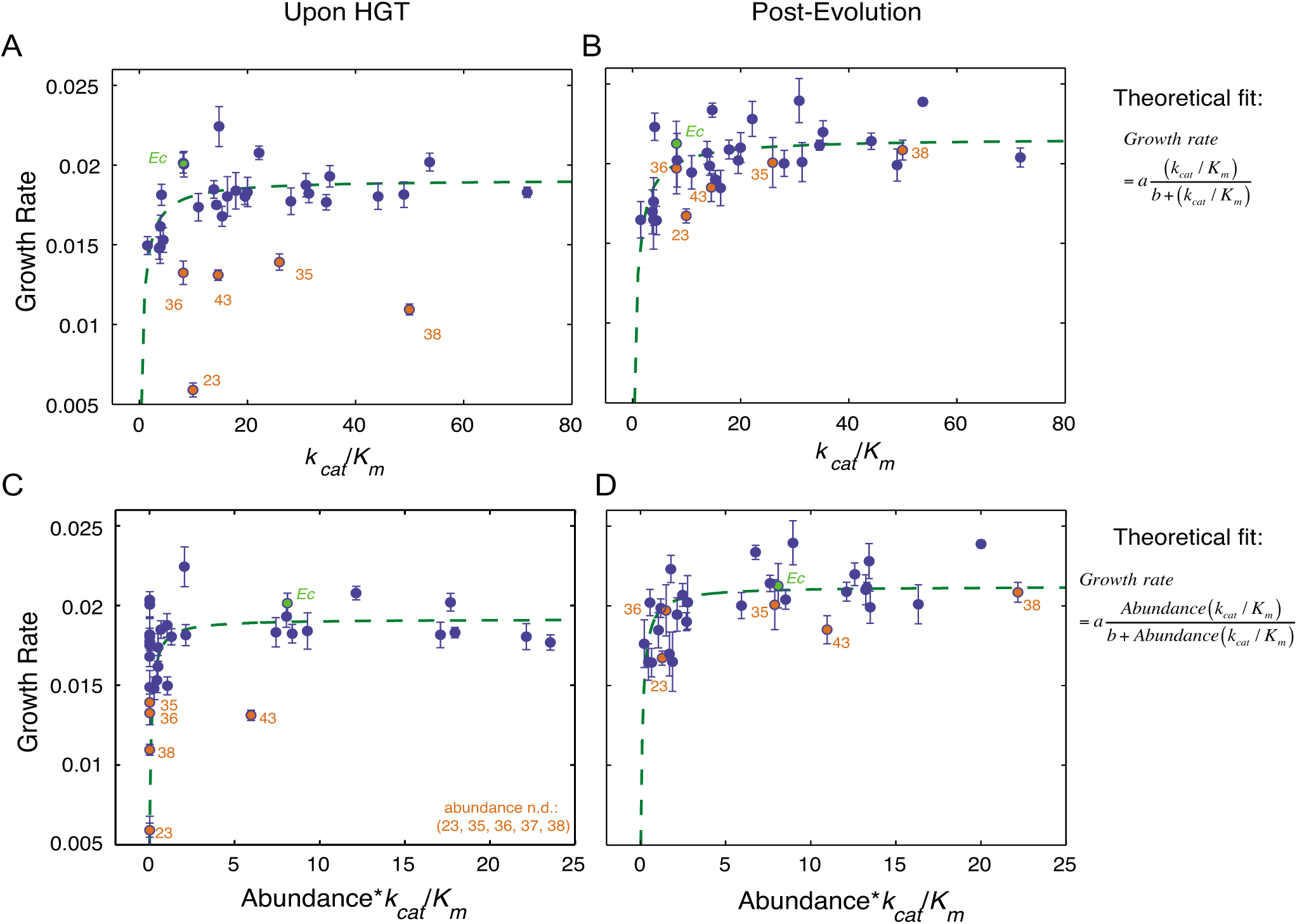
Fitness landscape of HGT strains is largely explained by flux dynamics theory. A) Relationship between activity (*k*_*cat*_/*K*_*m*_) of the purified orthologous DHFR proteins and growth rate of the naive HGT strains (prior to evolution experiment). Strains that exhibit the most dramatic drop in fitness (DHFR-23, 35, 36, 38, and 43) are highlighted in orange (see **Fig 1C**). Wild type *E. coli* is labeled *Ec*. No correlation between fitness and activity exists (R=0.34, p-value=10^−1^) when all points are included (see **Fig 4A**). However, after excluding the outliers (DHFR-23, 35, 36, 38, and 43), the relationship between activity and growth follows flux dynamics theory that predicts that growth rate is proportional to total folate turnover (see the equation on the right; constants *a* and *b* depend on the number of enzymes and topology of the metabolic network) (green line, p-value=10^−4^). B) Growth rate and activity after the evolution experiment. Similar to (A), fit also excludes DHFR-23, 35, 36, 38, and 43, but they nonetheless migrate to the prediction of the flux dynamics theory (n.d. – non-detectable). C-D) Relationship between growth rate and the product of relative intracellular abundance (from total lysates) and activity (*k*_*cat*_/*K*_*m*_). Theoretical fit also excludes DHFR-23, 35, 36, 38, and 43. (p-values are <10^−4^ for all theoretical fits in panels (A-D)).

Why did the activity-fitness relationship emerge only after experimental evolution? Catalytic activity alone does not fully account for the dependence of growth rates on DHFR properties because the effective functional level of intracellular DHFR is also a function of its intracellular abundance (i.e., the product [*abundance*\*(*k*_*cat*_/*K*_*m*_)]). Indeed, the mapping between dosage of an essential enzyme and fitness is predicted by flux dynamics theory [30]:

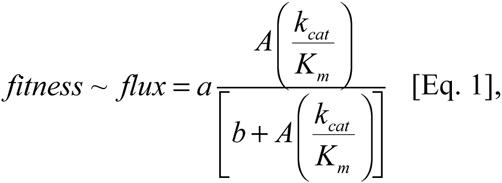

where *A* is the intracellular abundance, and constants *a* and *b* depend on the number of enzymes and topology of the metabolic network. We fitted this function for all strains, except the outliers (DHFR-23, 35, 36, 38, and 43). We found dosage-dependence not only for the evolved strains (**Fig 3D**), but also the naive strains that largely clustered around the theoretical fit (with the exception of DHFR-43 which remained an outlier) (**Fig 3C**). Consistent with the prediction from the flux dynamics theory (Eq. 1), strains with lower DHFR dosage than WT *E. coli* were less fit than WT *E. coli*, whereas strains with higher DHFR dosage did not necessarily appear more fit (plateau-like behavior) [31]. Additionally, the correlation between gene-dosage (the product [*abundance*\*(*k*_*cat*_/*K*_*m*_)]) and growth rate is statistically significant for both naive (Spearman r=0.56, p-value = 10^−3^) and evolved (Spearman r=0.64, p-value = 10^−4^) strains (**Fig 3C,D and 4A**). Thus, we conclude that the dosage-dependent fitness landscape of horizontally transferred DHFR genes generally accounts for the fitness effects of most HGT events. More generally, this mapping between fitness and the molecular properties of DHFR is an excellent manifestation of the “law of diminishing returns” [32,33]. Strains with *Abundance*\*(*k*_*cat*_/*K*_*m*_) equal or higher than WT generally lie in the neutral regime of the “plateau” and the rest are deleterious because of low *Abundance*\*(*k*_*cat*_/*K*_*m*_) values.

### Multi-dimensionality of the HGT fitness landscape

Next, we determined the cross-correlation among multiple parameters, including molecular and sequence properties of the orthologous DHFRs, their intracellular abundance, and fitness (**Fig 4A**, and **Materials and Methods**). As expected, there is no correlation between GC content and mRNA stability and growth rates **(Fig 4A**). Apart from the correlation of fitness with *Abundance*\*(*k*_*cat*_/*K*_*m*_) and (*k*_*cat*_/*K*_*m*_), which we noted is a consequence of the dosage-dependent fitness landscape (**Fig 3**), the only other significant correlations of fitness that we found was with promoter activity and charge (**Fig 4A-C**).

**Fig 4.**
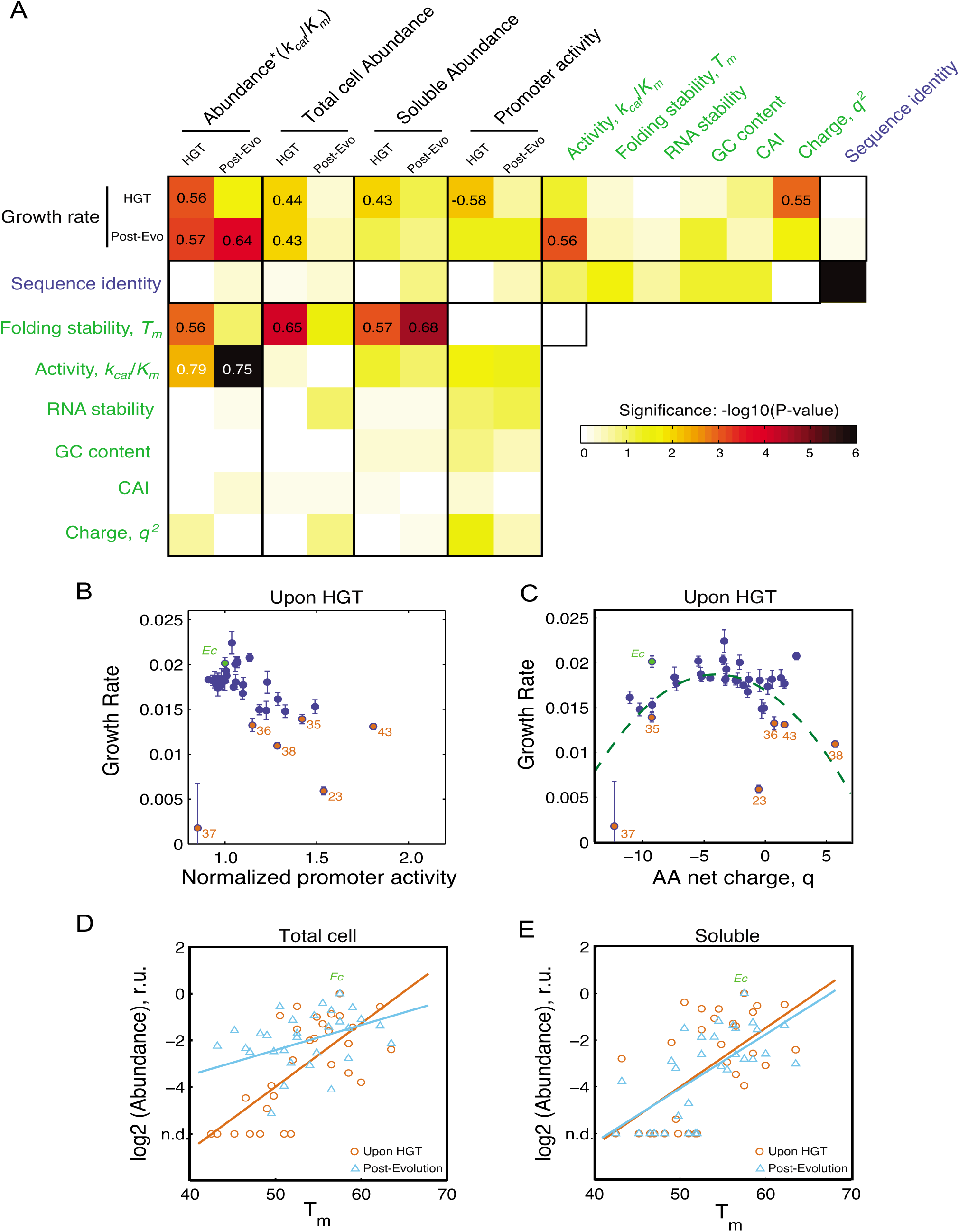
The role of molecular and cellular properties in the fitness effects of HGT. A) Correlation between growth rates, sequence and biophysical properties, and cellular responses (**S1-S3 Tables**) Blocks are colored according to –log10 (*p*-values) of Spearman correlation. Correlation coefficients with *p*-values<10^−2^ are indicated within the blocks. Promoter activity of the DHFR genes (endogenous *folA* promoter) is measured as a ratio between GFP signal from pUA66 plasmid and optical density of the growing cultures (see **Materials and Methods**). RNA stability is calculated over a 42 nt window starting near the translation site (-4) (see **Materials and Methods** and). Both RNA folding stability and GC content are calculated using the adapted DNA sequences (**S2 Table**). Neither GC content nor RNA folding stability correlates with fitness. Besides having a good correlation with k_cat_/K_M_ and abundance*k_cat_/K_M_ (**Fig 3**), fitness before evolution experiment shows significant correlation with promoter activity and net charge of DHFR. B) Activation of *folA* promoter is inversely correlated with growth rate before evolution, a signature of deficiency in intracellular DHFR abundance. This correlation disappears after the evolution experiment (**S4A Fig**), indicating an increase in DHFR abundance. C) Growth rates before evolution experiment exhibit an apparent quadratic dependence on the net charge *q* of the DHFR amino acid sequence. Dashed curve is the regression through the points defined by (q + 3.5)^2^ (p-value <10^−4^). The dependency becomes much less pronounced after the experimental evolution (**S4B Fig**). D,E) Intracellular DHFR abundance measured in total cell lysate (D) and soluble fraction (E) immediately before evolution correlates with folding stability *T*_*m*_ (orange circles) (r=0.65; p-value <10^4^ for total cell, and r=0.57; <10^3^ for soluble). After the evolution, a strong increase in DHFR abundance as measured in total cell lysates weakens the correlation with Tm (blue triangles; r=0.41, p-value=0.025) (D), but the strong correlation with Tm is preserved for soluble abundances (blue triangles; r=0.68, p-value<10^4^)

It was previously shown that when a cell loses the catalytic capacity of its DHFR, either through destabilizing mutations or treatment of the inhibitor trimethoprim (TMP), a feedback loop triggers up-regulation of *folk* promoter activity [34]. So we checked if the low abundance and lack of functional capacity upon HGT (**Fig 2C, Fig 3A,B**) could trigger the same behavior. Indeed, we found a highly statistically significant correlation between promoter activity and growth (orange dots in **Fig 4B**, r=-0.58, p-value=10^−4^). After experimental evolution ameliorates the deficiency in functional abundance (**Fig 3D**), the correlation between growth rate and promoter activity is lost (**Fig 4A** and **S4A Fig**). We note that the anti-correlation between promoter activity and fitness before evolution experiment is driven by the subset DHFR-23, 35, 36, 37, 38, and 43, which experience the most severe fitness loss. These results reaffirm that the direct consequence of the barrier is the drop in intracellular DHFR abundance.

Interestingly, we also observed a significant non-linear relationship (p<10^−3^) between the net charge (at pH 7) of the horizontally transferred DHFR proteins and growth rates of the HGT strains before experimental evolution (**Fig 4A,C** and **S1 Table**). This relationship follows a quadratic dependence, which reflects the observation that strains with reduced fitness also have DHFRs with high net charge. Again, the subset DHFR-23, 35, 36, 37, 38, and 43 drives the trend. The correlation of the DHFR with net charge is largely lost after the evolution experiment (**Fig 4A**, **S4B Fig**).

We also found a strong correlation between soluble and total intracellular abundances of DHFRs with thermodynamic stability in the strains upon HGT (p<10^−5^) (**Fig 4D,E**) (see **Materials and Methods**). Although this correlation remains strong for soluble abundance after the evolution experiment, this trend weakens for total abundance (**Fig 4A,D,E**). We found earlier, again in the context of mutations in DHFR, that the protein homeostasis degrades less stable proteins at a faster rate, thus giving rise to a relationship between *total* protein abundance and stability in *active cytoplasm*. In contrast, *in solution* stability affects only the distribution between folded and unfolded species leaving the total amount unaffected. Thus, the unchanged dependence of soluble abundance on stability after the evolution experiment is likely to be due to the *intrinsic* propensity of less stable proteins to aggregate, a property that has not changed throughout the evolution experiment. Altogether, these observations again point out to a potential role of protein homeostasis in HGT events.

### Lon protease is a major modulator of DHFR abundance upon HGT

Evidence suggest that loss of abundance is responsible for the barriers to horizontal transfer of the orthologous DHFR proteins, so next we seek to determine the molecular mechanism for that loss of abundance. To that end we performed whole genome sequencing (WGS) of the evolved DHFR-23, 35, 37, 38, and WT strains (**Fig 5A and S4 Table**). Out of 8 independent evolutionary trajectories conducted for each of the strains, trajectory 1 was arbitrarily chosen for sequencing. Randomly isolated individual clones as well as whole populations were sequenced for each of the strains (with the exception of DHFR-23, for which WGS was done only on the whole population, **S4 Table**) (see **Materials and Methods**). WGS revealed a number of genetic variations in the evolved strains at various allele frequencies, however, the *only* variation common to *all* genomes of the evolved HGT strains but *not* the evolved WT strain, was the IS186 insertion in the *clpX-lon* intergenic area (**Fig 5A and S4 Table).** The *lon* gene encodes ATP-dependent protease Lon that is central in maintaining protein homeostasis [35]. Lon is also known to directly control the turnover rates of several proteins in *E. coli* [36]. Previously, we showed that for *E. coli* strains carrying destabilizing mutations in the endogenous DHFR, *lon* deletion improved fitness by increasing the intracellular abundance of the mutant DHFR proteins. We surmised that a similar mechanism might have taken place in our HGT strains.

**Fig 5.**
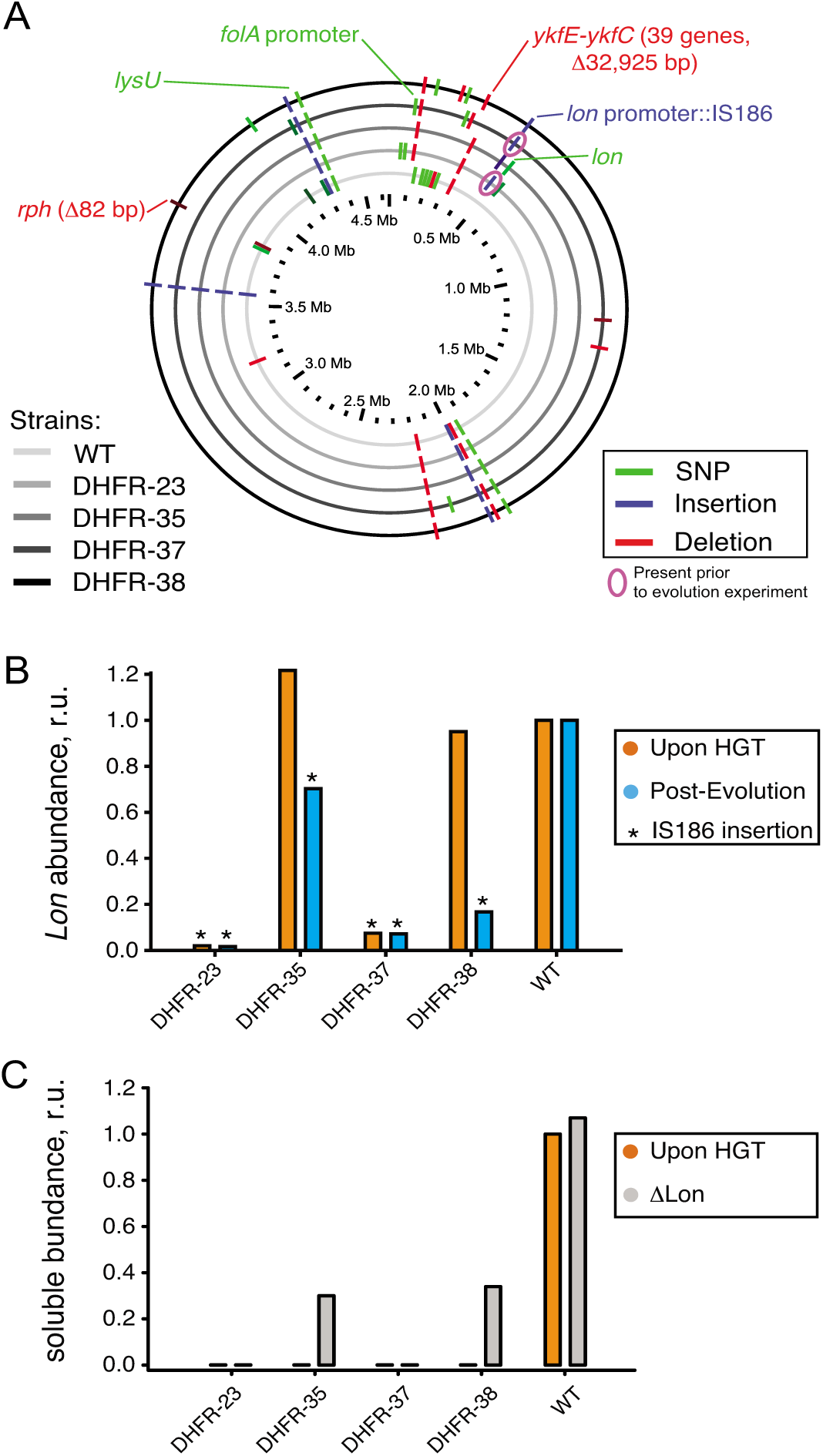
Sequencing of the orthologous strains. A) Mutations detected by whole genome sequencing (WGS) in the evolved populations of WT and orthologous DHFR-23, 35, 37, and 38 strains are indicated with respect to their location in *E. coli* chromosome. Only mutations that exceed 20% frequency in a population are shown (see **S4 Table** for detailed sequencing results). Mutations validated by PCR followed by Sanger sequencing (**S5 Table**), or known from literature are annotated. IS186 insertion in *clpX-lon* intergenic area was also found in naive DHFR-23 and 27 strains (*i.e.*, prior to evolutionary experiment) (pink circle). B) Intracellular Lon abundance decreases upon intergenic *clpX-lon* IS186 insertion. Intracellular Lon abundance in total cell lysates was detected using anti-Lon antibodies immediately upon HGT (orange) and after the evolutionary experiment (blue) (see **Materials and Methods**). Strains with IS186 insertion are marked with asterisk. C) Purifying *lon* knock-out (⊗*lon*) was performed on naive DHFR-23, 35, 37, and 38 strains (described previously in), and the resulted change in intracellular DHFR abundance in soluble fraction of cell lysates was measured with Western Blot using anti-His antibodies (see **Materials and Methods**). Abundance of DHFR-35 and 38 changed from an undetected levels to approximately 30% of the WT *E. coli* DHFR level.

To prove this conjecture, we randomly isolated 4-5 individual clones from the evolved populations of DHFR-35, 37, 38 and WT strains and performed a direct sequencing (PCR amplification followed by Sanger sequencing) of the *clpX-lon* intergenic area and *lon* coding sequence. We confirmed that the evolved HGT strains, but not the evolved WT, contain the IS186 insertion (see **Materials and Methods** and **S5 Table**). DHFR-35 exhibited a variety of outcomes whereby one colony contained the IS186 insertion in the *clpX-lon* intergenic area, one colony carried a non-synonymous D445E mutation in the *lon* coding sequence, and the remaining two colonies did not carry any mutations in the *lon* region (**S5 Table**). Next, we extended the direct sequencing of *clpX-lon* intergenic area to whole populations of *all* 35 naive and evolved HGT strains and WT strain (**S5 Table**). Again, only evolved strains of DHFR-23, 35, 37, 38 carried the IS186 insertion in the *clpX-lon* intergenic area, but not the evolved WT strain. This same insertion was detected earlier in bacterial populations that developed antibiotic resistance and was hypothesized to inactivate Lon by blocking its expression [37], although no experimental evidence was provided to support it. We tested this hypothesis in the current work by measuring Lon intracellular abundance (see **Materials and Methods**) and found that all the HGT strains carrying the IS186 insertion upstream to their *lon* gene have indeed dramatically reduced the intracellular Lon abundances compared to WT (**Fig 5B**). These are also the strains that exhibited a dramatic improvement in their fitness upon conclusion of the evolutionary experiment (**Fig 1C**).

Interestingly, we also found that DHFR-23 and 37 carried the IS186 insertion prior to the evolutionary experiment, whereas other naive HGT strains and naive WT did not (**S5 Table**). The WT *E. coli* strain (MG1655) used in our experiment to generate the HGT strains also does not carry the IS186 insertion in the *clpX-lon* intergenic area as shown by direct PCR sequencing.

By virtue of the experimental design, each HGT strain in our experiments originated from a single WT cell where a homologous recombination successfully replaced the endogenous *folA* gene with a DHFR ortholog (see **Materials and Methods**). However, because of the heavy fitness cost associated with orthologous replacements of the endogenous *folA* gene with genes encoding DHFR-23 and 37, we believe that a homologous recombination only resulted in a viable strain if it was accompanied by a compensatory mutation. It is plausible that incorporation of IS186 transposable element in the *lon* promoter, a known hot spot for IS186 incorporation [38], was co-selected with the DHFR-23 and 37 replacement because it buffered the otherwise severely deleterious (possibly lethal) fitness effects. For DHFR-35 and 38, the IS186 insertion in the *clpX-lon* intergenic region was not present in the naive strain, but only arose as a result of the experimental evolution, which was accompanied by a dramatic improvement in fitness. Altogether, the IS186 insertion in the *clpX-lon* intergenic region was selected *independently* in four different HGT lineages: Twice as an immediate compensation of the deleterious fitness cost associated with orthologous replacement (naive DHFR-23 and 37 strains), and twice as a result of experimental evolution (evolved DHFR-35 and 38 strains).

Since we observed *lon* inactivation via IS186 insertion only in evolved orthologous transfer strains, *but not evolved WT*, the direct test of whether Lon is solely responsible for the phenotype would be to knock out *lon* in the naive orthologous transfer strains, and follow the ensuing effect on fitness and DHFR abundance. To that end we introduced *lon* knock-out mutations on the background of naive (pre-experimental evolution) HGT strains. Previously, we observed that *lon* knock-out resulted in substantial improvement in growth of most naive HGT strains, including DHFR-23, 35, 37 and 38. In the current work, we measured the effect of *lon* knock-out on the intracellular DHFR abundance. We indeed found a marked increase in abundances of DHFRs 35 and 38 in a soluble lysate fraction (an increase from non-detectable levels to levels that comprise ∼30% of abundance measured for *E. coli* DHFR in WT strain) (Fig. 5C). Thus, the underlying mechanism behind the detrimental fitness effects of HGT is indeed the action of Lon on the foreign DHFR.

### Gene-specific mechanism to fine-tune DHFR abundance follows Lon inactivation

Despite the strong selective advantage of shutting down the Lon protease, especially in the background of very deleterious HGT events (**Fig 4**), such a mechanism may not be evolutionary sustainable because Lon has other clients that are central to cellular function [36]. Thus, we hypothesized that mutations specific to DHFR would eventually arise to maintain its required intracellular abundance. This may include mutations in the DHFR promoter and/or coding regions, or gene duplication events.

To explore this possibility, we investigated in greater detail 8 independent evolutionary trajectories of DHFR-37, which suffered a deleterious effect from HGT but dramatically increased fitness after the evolution experiment in all trajectories (**Fig 6A**). Indeed, whole-genome sequencing of the evolved DHFR-37 (a single evolutionary trajectory randomly chosen out of 8 independent trajectories after 31 serial passages) revealed a C to T mutation at position - 35 of the *folA* promoter with a concomitant increase in the intracellular DHFR abundance (**Fig 6B**). Interestingly, this mutation was previously shown to increase the trimethoprim resistance in *E. coli* MG1655 strain, presumably by increasing DHFR abundance [39]. As noted in the previous section, DHFR-37 had the IS186 insertion inactivating the activity of its Lon protease already at the naive stage (**Fig 5**).

**Fig 6.**
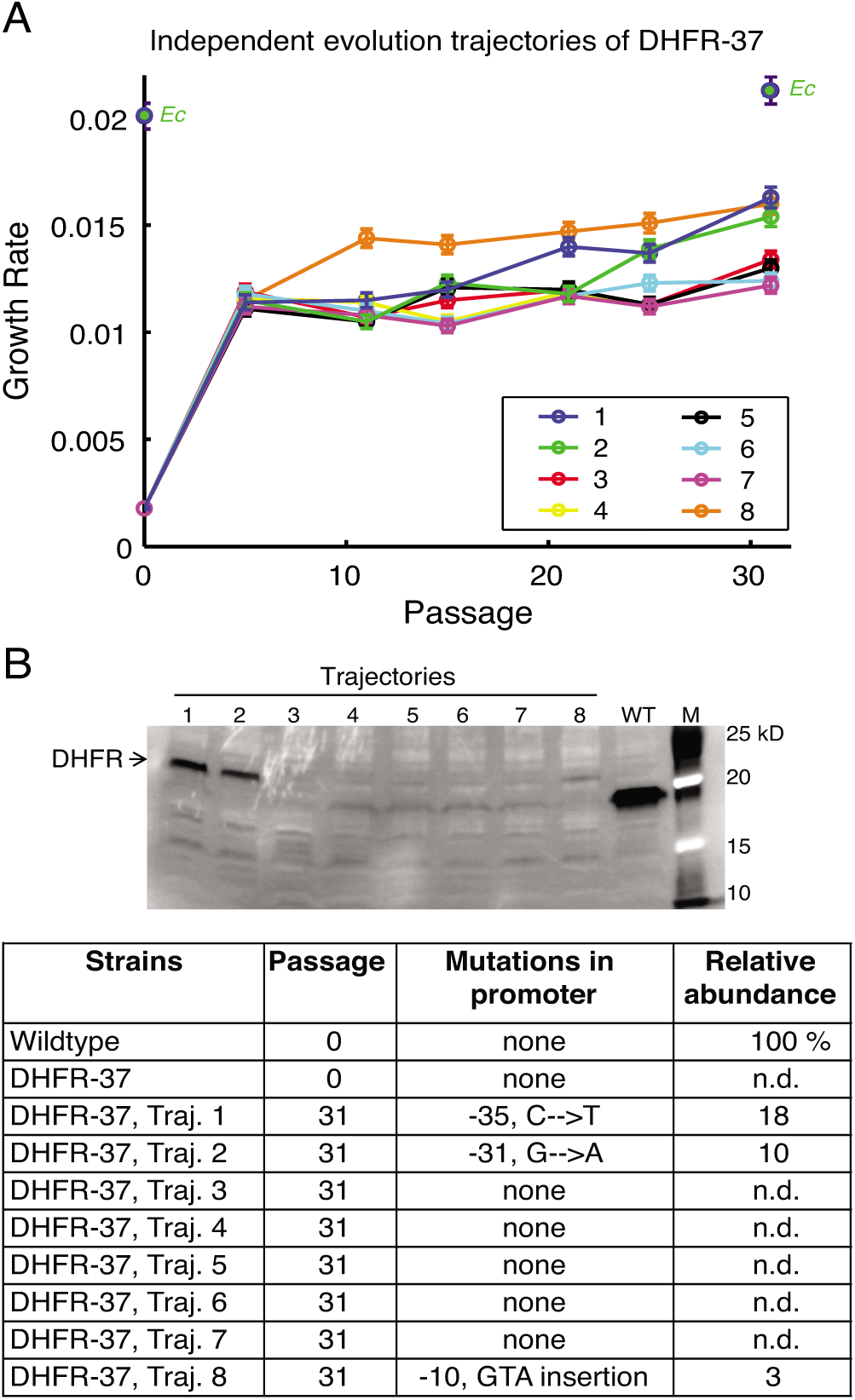
Fine-tuning evolution of orthologous DHFR expression. A) Eight independent evolutionary trajectories of DHFR-37 (from *P. ananatis*) show an increase in fitness after evolution. Growth rates of each individual trajectories were measured every 5 passages. Three trajectories (1, 2, and 8) become markedly fitter than the rest after 31 passages. Error bars represent standard deviation of 4 independent measurements. B) Soluble abundance of DHFR-37 protein was measured in soluble cell fractions of all eight trajectories after evolution (In the gel, M: Marker). Note the pronounced abundance levels in trajectories 1, 2, and 8. N.d. not detected. Sequencing of *folA* gene revealed characteristic mutations in the promoter region that explain the increased abundance and improved fitness.

We performed direct sequencing of the *folA* gene in four randomly chosen colonies from each of the 8 independent evolutionary trajectories of DHFR-37 strains. As shown in **Fig 6B**, we found that two more trajectories acquired mutations in *folA* promoter: G to A at position -31, (trajectory 2) and a GTA insertion at position -10 (trajectory 8). Similarly to trajectory 1 (C to T substitution at -35), these mutations were accompanied by an increase in DHFR abundance (**Fig 6B**) and improvement in growth rate (**Fig 6A**). Thus, these results show that once the barrier imposed by the proteostasis is overcome, a more fine-tuned evolutionary response follows.

### Proteome-level responses upon HGT and after crossing the fitness barrier

To determine how the effect of HGT percolates throughout the entire *E. coli* proteome, we next analyzed the systems-level effect of inter-species DHFR replacements before and after the evolution experiment. To that end, we quantified relative (to WT) abundances of ∼2000 proteins in the cytoplasm using tandem mass tags (TMT) with subsequent LC-MS/MS analysis (**Materials and Methods** and [34]). We picked five strains for proteomic characterization based on their fitness effect upon HGT (**Fig 1B**): DHFR-23, 35 and 38 (severely deleterious); DHFR-22 (mildly deleterious); and DHFR-39 (beneficial). For reference, we compared the proteomic effects of orthologous replacements with the proteomic effect of treating *E. coli* with 1 |g/ml of trimethoprim (TMP). For each of the >2000 proteins detected, we quantified their enrichment using the log of relative protein abundances (LRPA) that are expressed as z-scores (see **Materials and Methods**). In **Fig 7A**, we show the correlation plots of the z-scores between proteomes. The complete gene-by-gene proteomics data are presented in **S6 Table**.

**Fig 7.**
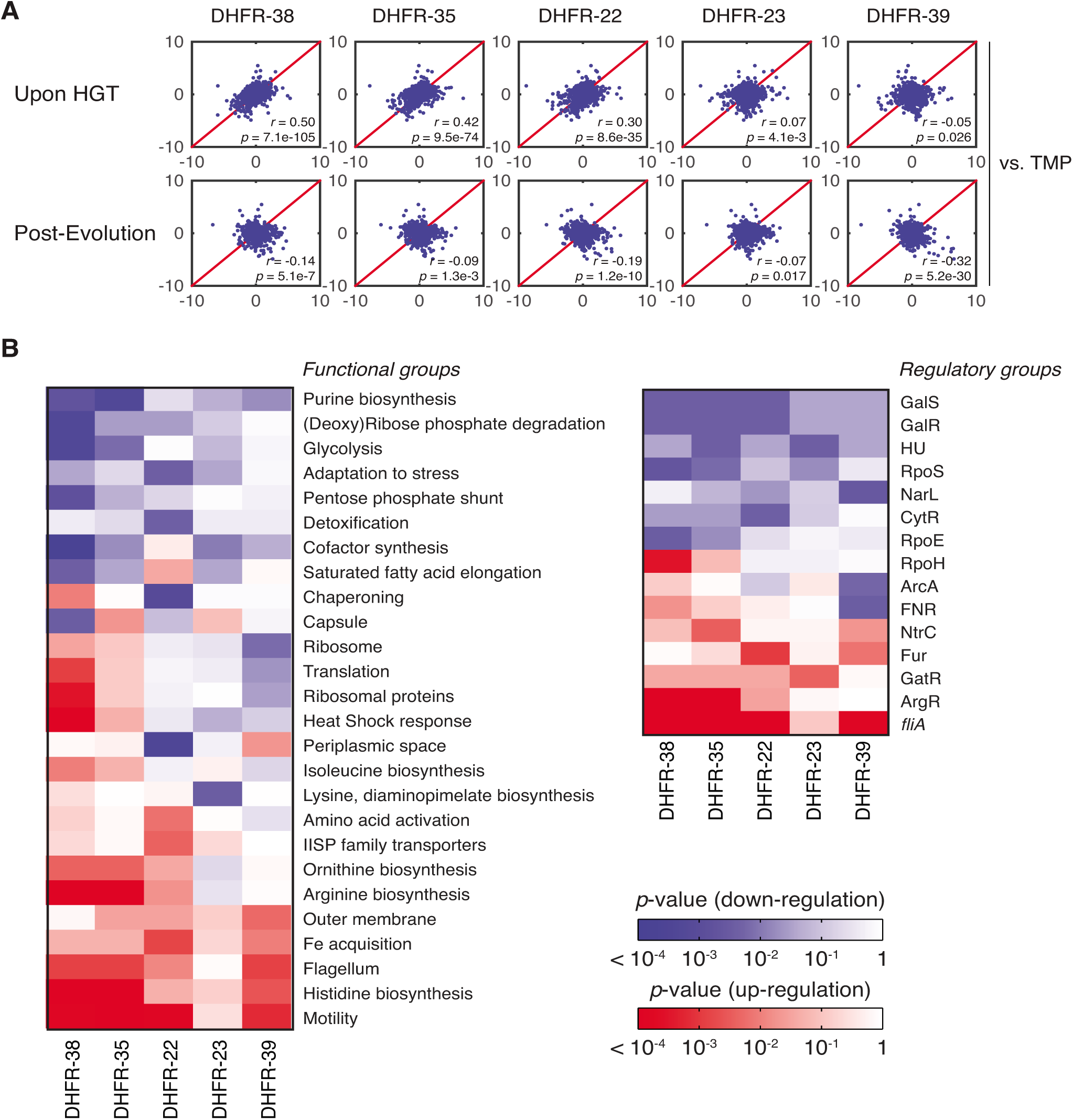
Global proteomic response to HGT. A) z-scores correlation plots between proteomes of indicated DHFR orthologous strains (upon HGT and after evolution) and WT strain treated with 1 |g/ml trimethoprim (TMP) (see **Materials and Methods** and **S6 Table**). The strains are representative of the fitness effects upon HGT: DHFR-23, 35 and 38 are severely deleterious; DHFR-22 is mildly deleterious; and DHFR-39 is beneficial (**Fig 1C**). Global proteome quantification was obtained using LC-MS/MS analysis of TMT-labeled proteomes [34]. Each quantified protein (+2,000 proteins per proteome) is assigned a z-score that measures the enrichment of its abundance relative to the whole proteome (see **Materials and Methods**). The proteome profile upon HGT of DHFR-38, 35 and 22 is similar to the adverse cellular state during the antibiotic treatment, but is relaxed after experimental evolution. The strains are ordered left to right according to decreasing similarity with TMP treated WT cells (top panel). B) Change in global variation in protein abundances induced by experimental evolution of the indicated orthologous strains calculated for functional and regulatory classes of genes. Color code indicates logarithm of p-values of two-sample KS tests performed on pre-and post-evolution proteomes along with the direction of change (blue – drop in abundance, red – increase in abundance) (see **Materials and Methods**). We plotted only the gene groups whose change upon evolution experiment is significant (KS p value less than 0.01). The proteomic responses DHFR-35 and 38 upon HGT and post-evolution are similar despite their evolutionary distance (see **Fig 1C**).

As expected for DHFR-35 and 38, where HGT is severely deleterious, their proteomes strongly resemble the proteome of TMP-treated wild-type strain (*r* = 0.42, *p*-value = 9.5×10^−74^ and *r* = 0.50, *p*-value = 7.1×10^−105^, respectively) (**Fig 7A**). This result suggests that the systems-level response to HGT of the DHFR genes is akin to response to inactivation of the endogenous DHFR protein by TMP. Interestingly, despite significant evolutionary distance between the DHFR alleles from strains DHFR-35 and 38 (**Fig 1C**), the correlation between their proteomic profiles is significant (*r* = 0.84, *p*-value < 10^−300^). However, the proteome of DHFR-23, another orthologous strain with a severely reduced growth, was not similar to the TMP-treated WT proteome (*r* = 0.07, *p*-value = 0.0041), suggesting that, at least for some strains, the systems-level response to the partial loss of DHFR function follows a different pattern. The proteome of DHFR-22, a strain with moderately reduced fitness, was much less similar to the proteome of TMP-treated WT (*r* = 0.30, *p*-value = 8.6×10^−35^). The proteome of DHFR-39, one of the few strains that grew better than WT upon HGT, bore no resemblance to TMP treatment (*r* = -0.05, *p*-value = 0.026).

After the evolution experiment, the proteomic profiles of the strains lose their resemblance to TMP-treated WT cells (**Fig 7A**), which reflects the alleviation of the detrimental effects of HGT (**Fig 3B,D**). Additionally, after the evolution experiment the proteomic profiles of the strains become more similar (see **S6 Fig** and note the increased correlation in inter-strain comparison). In particular, although the proteome of DHFR-23 is barely similar to DHFR-35 and 38 before the evolution experiment, it becomes similar to DHFR-35 and 38 after it (**S6 Fig**). This observation is consistent with the clustering of the strains near the growth rate-metabolic turnover curve that is predicted by flux dynamics analysis (**Fig 3B,D**).

Next, we carried out a comparative analysis at the level of functional pathways and operons. Using the functional and regulatory classification of genes by Khodursky and co-workers [40], we screened the gene groups that collectively changed their abundances significantly during the evolution experiment (**Fig 7B**), using the Kolmogorov-Smirnov test (see **Materials and Methods**). Among them we found that the genes responsible for cell motility show the largest increase in their abundances, which is notable, because it was previously reported that the loss of DHFR function either by TMP treatment or destabilizing mutations causes a sharp down-regulation of the most of the motility genes [34]. This suggests that through the adaptation process, HGT strains have eliminated the energetic burden which caused shutting down of cell motility in the first place (fliA operon and motility class in **Fig 7B**). In the same vein, genes responsible for a number of metabolic processes such as synthesis of amino acids and nucleotides as well as turnover of several metals show highly significant changes between the naive and evolved strains (**Fig 7B**).

## DISCUSSION

In this work we provided molecular-level insight into the origin of barriers that shape the fitness landscape of HGT events. We focused here on xenologous transfer of DHFR coding genes, a common mode of HGT [1]. The chromosomal gene replacements were done while preserving the endogenous promoter. Such an experimental design provided us with a direct control over conditions of expression, enabling us to focus on the link between variation of sequence and biophysical properties of transferred proteins and the ensuing fitness effects of HGT. We were able to purify 33 orthologous DHFRs and measure their activities and stabilities. Over half of them are more catalytically active and stable than *E. coli* DHFR. Nevertheless, inter-species replacement of *E. coli* DHFR with non-toxic and much more catalytically active proteins resulted in a considerable loss of fitness.

Experimental evolution revealed the main culprit of the severe loss of fitness – Lon protease, which is repeatedly deactivated in strains with most dramatic fitness improvement. While previous work showed that *lon* knockout could rescue fitness of *E. coli* containing orthologous DHFRs (since Lon protease is crucial for DHFR turnover), over-expression of the chaperonins GroEL/ES produced a similar fitness improvement [17]. However, we show here that actual evolutionary processes consistently utilize Lon deactivation, rather than GroEL/ES over-expression. Given their similar fitness effects, this suggests that Lon deactivation must be more accessible by the processes that produce the genetic variation on which selection acts. Indeed, we find that Lon deactivation is consistently achieved by insertion of mobile elements in the *lon* promoter; mobile element insertion is known to occur frequently in bacterial genomes and thus is a major source of genetic variation [27].

Is the partial shutdown of the degradation branch of proteostasis a generic response to HGT events? Lon inactivation increases the number of functional enzymes, but it also increases the number of misfolded/aggregated species that can be toxic in highly abundant proteins [41]. The toxicity due to misfolding may be less relevant for HGT events involving low abundance proteins. The DHFR abundance in *E. coli* is about 40 molecules per cell [24], on the lower end of the range of abundances for the rest of the *E. coli* proteome, which span at least five orders of magnitude (10^−1^ to 10^4^ protein copies per cell in single-cell measurements [24] or 10^1^ to 10^6^ per cell in bulk measurements [42]). Our work unambiguously demonstrates the role of proteostasis in shaping HGT fitness. However, determining the specific response of the proteostasis machinery to different HGT events is an exciting subject of future research.

The key finding of this work is that *E. coli* cells apparently discriminate between own and foreign proteins. Most likely the discrimination is post-translational: the *folA* promoter gets activated as cells lose DHFR function, similarly to the effect of DHFR point mutations and inhibition by TMP [34], so that the loss of the protein is due to post-translational degradation and/or aggregation.

The mechanism of recognition of foreign proteins by *E. coli* proteostasis machinery remains somewhat of a mystery. In earlier studies Lon was implicated in degradation of destabilized DHFR and other misfolded proteins [43]. However in this case orthologous DHFRs do not appear unfolded or misfolded, at least in solution in vitro. Global molecular properties of orthologous DHFRs such as amino acid composition or net charge do not correlate with their abundances in *E. coli.* We also do not observe a correlation between the evolutionary distance of orthologous DHFRs from *E. coli* and fitness of the HGT strains. It is for the future studies to establish sequence or structural/physical signatures that trigger degradation of these proteins in *E. coli* cytoplasm.

Current phylogenetic methods can now detect HGT events in a wide range of evolutionary ages, even within pathogenic clone outbreaks [44]. However, these bioinformatics approaches are agnostic about molecular and cellular mechanisms that incur fitness costs or benefits of HGT events. As shown in this work, pleiotropy at the level of molecular properties and cellular components could shape the HGT fitness landscape. Swapping out a protein with an orthologous copy (or adding point mutations to a WT protein [45]) has a much broader effect than merely changing the protein’s individual molecular properties like stability and activity [34]. As we observe here, HGT events that actually improve DHFR activity may nevertheless be deleterious at first because they interact negatively with proteostasis machinery of the host. We believe that experimental studies to dissect the mechanistic origin of pleiotropy will complement our understanding of HGT from genomics by pointing out to other molecular or sequence signatures of HGT events.

Our finding that elements of the proteostasis machinery have the ability to discriminate between “self” and “non-self” proteins points to an important possible constraint on evolution of proteomes to maintain compatibility with their own proteostatic machinery. Future studies might reveal the manifestations of this evolutionary constraint, such as certain proteases and chaperones evolving slower than other components of proteomes, or co-evolution of sequences of components of proteostasis machinery with global features of the whole proteome such as charge distributions or amino acid compositions.

Besides being relevant to understanding the evolutionary dynamics of HGT, our approach is broadly applicable to the study of the genotype-phenotype relationship. While the concept of a fitness landscapeis dominant in evolutionary biology, it remains highly metaphoric as its “axes” remain unlabeled. A promising approach to map fitness landscape is by introducing ‘’bottom up,’’ controllable genomic variations that cause known changes of the molecular properties of proteins [17,45-51]. However, point mutations and/or random mutagenesis are limited in their ability to generate a broad variation of catalytic activity and other physical properties of proteins. But “borrowing’’ highly diverged yet catalytically active orthologous proteins from other species allows us to cover a broad range of variation of molecular properties of proteins. By systematically exploring the relation between molecular properties of xenologously replaced proteins and fitness this approach provides an opportunity to quantitatively characterize the global properties of fitness landscapes.

## Materials and Methods

### Orthologous DHFR sequences and cloning

The BLAST analysis against E. *coli*’s DHFR amino acids sequence of mesophilic bacteria produced 290 unique DHFR sequences [52]. This dataset was used to select 35 sequences with amino acid identity to E. *coli*’s DHFR ranging from 29 to 96% (**Fig 1**, **S1 Table**). The amino acid sequence of the chosen DHFRs was converted into the DNA sequence using the codon signature of the E. *coli*’s *folA* gene. Specifically, the frequency of each codon was calculated, and codons with the highest score were used for protein – DNA sequence conversion (**S2 Table**). In addition, each DNA sequence, including E. *coli*’s *folA*, was fused to a tag encoding 6 histidines. The resulted DNA sequences were synthesized (GenScript) and cloned into pET24a+ plasmid (EMD Millipore) for recombinant expression and purification, and finally cloned into pKD13 plasmid for homologous recombination (see below).

### Homologous recombination

The detailed description of the method can be found in. Briefly, orthologous DHFR sequences placed under the *E. coli’s folA* endogenous regulatory region (191 bp separating the stop codon of the upstream *kefC* gene and the start codon of *folA* gene) were ligated into a pKD13 plasmid flanked by two different antibiotic markers (genes encoding for kanamycin (kanR) and chloramphenicol (cmR) resistances). The entire cassette was then amplified with two primers tailed with 50 nucleotides homologous to the chromosomal region intended for recombination (*kefC* gene upstream and *apaH* gene downstream of *folA)*. The amplified product was transformed into BW25113 strain with induced Red helper plasmids, and the recombinants selected on plates carrying both antibiotics. Strains carrying the putative orthologous replacement in the chromosome were verified by sequencing. Identified chromosomal replacements were then moved to MG1655 strain by P1 transduction and double antibiotic selection (kan and cm) and again verified by sequencing. Absence of the endogenous *E. coli’s folA* gene lingering in the orthologs strains was verified by PCR and by Western Blot with specific anti-*E.coli*’s DHFR antibodies.

### Media and growth conditions

Cells were grown from a single colony overnight at 30°C in M9 minimal medium supplemented with 0.2% glucose, 1 mM MgSO4, 0.1% casamino acids, and 0.5 |g/|L thiamine. Overnight cultures were diluted 1/100 and grown at 37°C. Growth rate measurements were conducted for 10 hours. OD data were collected at 600nm at 20 min intervals. The resulting growth curves were fit to a bacterial growth model to obtain growth rate parameters [53].

### Laboratory evolution

Strains were propagated in deep 96-well plates in 8 independent trajectories by serial passaging from a diluted culture to saturation in 12-hour growth cycles at 37°C using the TECAN robotic liquid handling system. We performed 31 of such passages which amounts to ∼600 generations (for the wild-type strain), assuming a continuous exponential phase.

### mRNA folding stability and codon adaptation index

mRNA folding stability was calculated using the HYBRID-SS-MIN routine of mFold over a window of 42nt starting with -50 upstream of the ATG of the DHFR coding region using the default parameters (http://mfold.rna.albany.edu/?q=unafold-man-pages/hybrid-ss-min) [54]. Codon adaptation index (CAI) was calculated as previously reported [55] using the whole *E. coli* genome as the reference set.

### Protein purification

pET24 expression vectors carrying the orthologous DHFR sequences fused to C-terminal (6x)His-tag under isopropyl β-D-1-thiogalactopyranoside (IPTG) inducible T7 promoter were transformed into BL21(DE3) cells. Cultures were grown at 30°C overnight from a single colony in Luria broth (LB) supplemented with 2% glucose, diluted 1/100 into Terrific Broth (TB) and grown for 4 hours at 28°C. Cultures were then chilled to 18°C, supplemented with IPTG (0.4 mM final concentration) and grown overnight. The recombinant proteins were purified from a lysate on Ni-NTA columns (Qiagen) followed by gel filtration to over 95% purity.

### Enzyme kinetics

DHFR kinetic parameters were measured by progress-curve kinetics, as described in (Fierke et al., 1987). Purified enzymes (10 nM) were pre-incubated with 120 |M NADPH in MTEN buffer (50 mM 2-(N-morpholino)ethanesulfonic acid, 25 mM tris(hydroxymethyl)aminomethane, 25mM ethanolamine, and 100 mM sodium chloride, pH7). Reaction was initiated by addition of dihydropholate (20, 15, 10 |M final concentration) and monitored till completion at 25°C following the drop in absorbance signal at 340 nm (NADPH disappearance) in Carry 60 spectrophotometer (Agilent). The kinetics parameters (k_*cat*_ and K_m_) were derived from progress-curves analysis using Global Kinetic explorer (Johnson et al., 2009).

### Stability measurements

Thermal stability was characterized by Differential Scanning Calorimetry (DSC), essentially as described [56]. Briefly, DHFR proteins in Buffer A (10 mM potassium-phosphate buffer pH8.2 supplemented with 0.2 mM EDTA and 1 mM beta-mercaptoethanol) were subjected to temperature increase at a 1°C/min rate between 20 to 80°C (nano-DSC, TA instruments), and the evolution of heat was recorded as a differential power between reference (buffer A) and sample (40 |M protein in buffer A) cells. The resulting thermograms (after buffer subtraction) were used to derive apparent thermal transition midpoints (T_m_ ^app^) from the peak in heat consumption.

### Plasmid transformation with WT and orthologous DHFRs

Genes encoding the orthologous DNA sequences were cloned into pBAD expression vector (EMBL) under the control of arabinose inducible promoter, and transformed into wild-type *E. coli* strain. For complementation assay, pBAD plasmid carrying *E. coli*’s DHFR (WT) was transformed into strains carrying orthologous replacements of the *folA* gene. The transformants were grown from a single colony overnight at 30°C in the supplemented M9 medium (+ 100 |g/ml amplicillin), diluted 1/100 and grown at 37°C in presence of 0.01% arabinose to achieve 50-100 fold increase in expression, relatively to the baseline chromosomal expression (for pBAD-phylogeny DHFR), or without the inducer to achieve 6-8 fold increase in expression, relatively to the baseline chromosomal expression (for pBAD-wtDHFR) (see **S2B Fig**). Growth rates were determined as described above.

### Promoter activity

Strains were transformed with pUA66 plasmid carrying *folA* promoter fused to GFP coding gene (Zaslaver et al., 2006). Promoter activity is defined as a ratio between fluorescent signal (excitation 495 nm, emission 510 nm) and biomass production (measured as OD at 600nm).

### Intracellular protein abundance

Cells were grown in supplemented M9 medium for 4 hours at 37°C, chilled on ice for 30 min and lysed with BugBuster (EMD Millipore). DHFR amounts in the soluble fraction were determined by SDS-PAGE followed by Western Blot using mouse anti-His-Tag monoclonal antibodies (Rockland Immunochemicals) and goat anti-mouse polyclonal secondary antibodies conjugated with WesternDot 625 (Life Technologies). Total Lon amounts were quantified using anti-Lon antibodies (a generous gift from the T. Baker’s lab) following the procedure described in [57]

### Whole-genome-sequencing

Genomic DNA was extracted using E.Z.N.A Bacterial DNA kit (Omega Bio-Tek) following the manufacturer’s instruction. Sequencing was performed on whole-population samples on Illumina MiSeq in 2x150 bp paired-end configuration (Genewiz, Inc., South Plainfield, NJ). Sequencing was also performed on samples derived from an individual colony from one evolutionary trajectory (out of eight). The raw data were processed with the *breseq* pipeline [58] on default settings, using the *E. coli* K-12 MG1655 reference genome (GenBank accession no. U00096.3) with a modified *folA* locus for each strain. Average coverage was ∼100-200 bases for each sample. Only mutations with >20% frequency were considered. We rejected variants that only appeared on reads aligning to a single strand as well as variants in loci with significant alignment problems. We confirmed all variants using a separate alignment method (BWA (http://arxiv.org/abs/1303.3997v2) and SAMtools [59]) and manually inspected each variant using IGV [60]. The results are summarized in S4 Table.

### Direct sequencing

To validate WGS data, we designed specific primers to amplify the genomic alleles of interest either from whole populations or independent colonies. Primers GGTGCGTTTGCCGGTCTGGATAAAGTG (forward primer) and CGGTGCCGTCAGGCAGTTTCAGCATC (reverse primer) were used to amplify the ClpX-Lon intergenic region, while primers GTGAAGCACAGTCGTGTCATCTG (forward) and CACTTGAATCCTTCAAGGTACGAACGCG (reverse) were used to amplify the lon ORF. The PCR products were subsequently sequenced by Sanger method.

### Proteomics

For global proteome analysis, whole populations of the evolved strains cells were lysed into 50 mM NaH_2_PO_4_ buffer (pH8) supplemented with BugBuster extraction reagent and benzonase (EMD Millipore), and soluble fraction was separated by centrifugation. Soluble cell lysates were trypsinized overnight by Promega (Madison, WI) Trypsin/Lys-C enzyme mixture with ratio 1:30 enzyme to protein and labeled with TMT reagent (TMT^®^, Thermo, San Jose, CA) followed by nano LC-MS/MS separation and analysis (for detailed description of the method see [34]). Z-scores of the log of relative (to wild-type) protein abundance (LRPA) were obtained according to Eq (1):

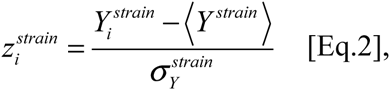

where index *i* refers to gene, 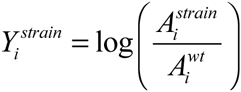 is LRPA for gene *i*, 〈*Y ^strain^* 〉denotes an average quantity *Y*_*i*_ over all genes for a given strain or condition in corresponding experiments, and 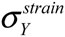 is a standard deviation of 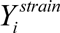.

### Kolmogorov-Smirnov Test on Functional and Regulatory Gene Groups derived from LC-MS/MS analysis

After grouping genes according to Sanguardekar et al. [40], we employed the Kolmogorov-Smirnov test in order to check how much a certain gene group changed upon serial passaging. First, we determined the direction of adaptation by comparing the average z-values of the naive and evolved proteome sets of a given gene group. If the average z-value increases, it is considered up-regulation; otherwise, it is considered down-regulation. Then, we applied the two-sample Kolmogorov-Smirnov test on the two sets, which provides the p-value for the null hypothesis that the two sets were drawn from the same distribution. Hence, we can quantitatively interpret a lower p-value as an indication that the two sets have more different distributions of z-values.

## Acknowledgements

We thank Daniel Hartl and members of the Shakhnovich lab for useful discussions.

## Supporting Information Figures and Legends

**S1 Fig.**
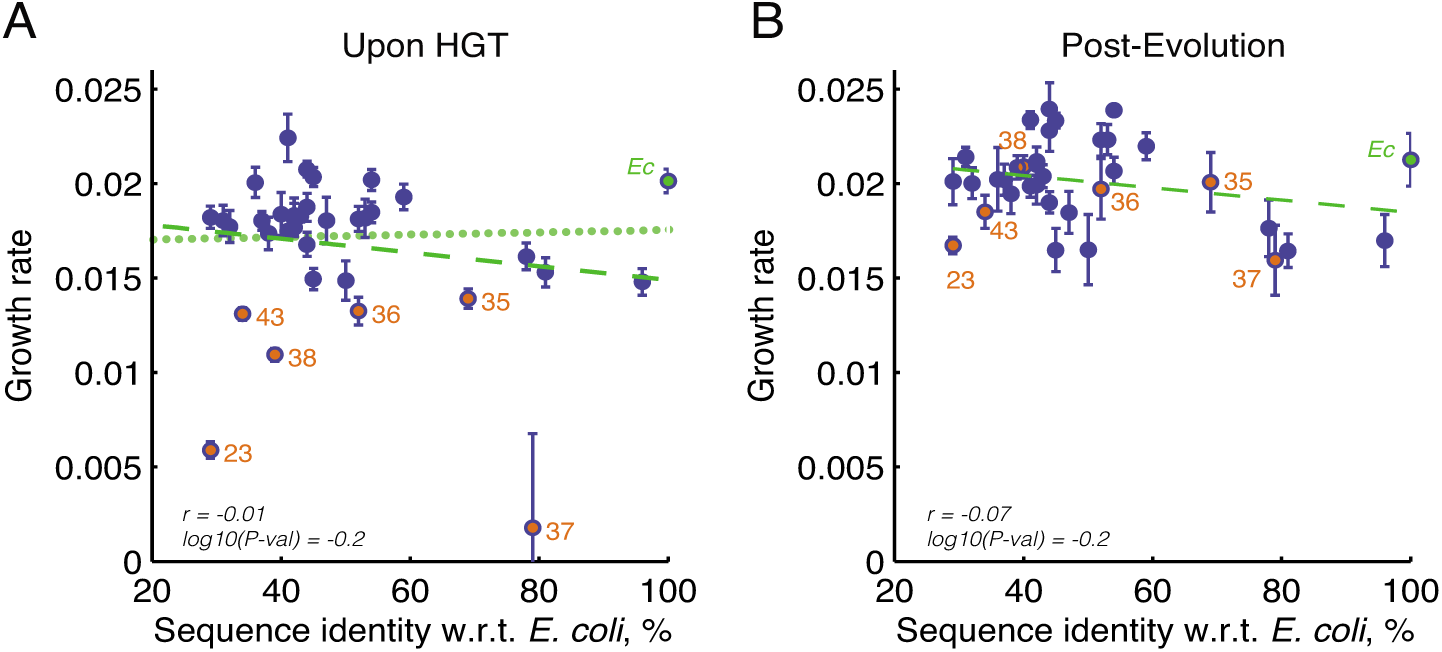
Evolutionary distance of orthologous DHFR proteins does not correlate with fitness effects of HGT. A,B) Evolutionary distance of orthologous DHFRs, measured as % of amino acids sequence identity with respect to WT *E. coli* DHFR (see **S1 Table**). Sequence identity does not correlate with growth rates before (A) or after (B) evolution (Spearman r and p-values are indicated). *Ec* (green) denotes WT *E. coli* strain. Strains with severe fitness effects are highlighted in orange. Dashed green lines are regression fits to all points. Excluding DHFR-37 in (A), there is still no correlation between identify and growth rate (r=0.06, p-value=0.72; dotted line is regression fit excluding DHFR-37). Error bars represent standard deviation of 4 independent measurements.

**S2 Fig.**
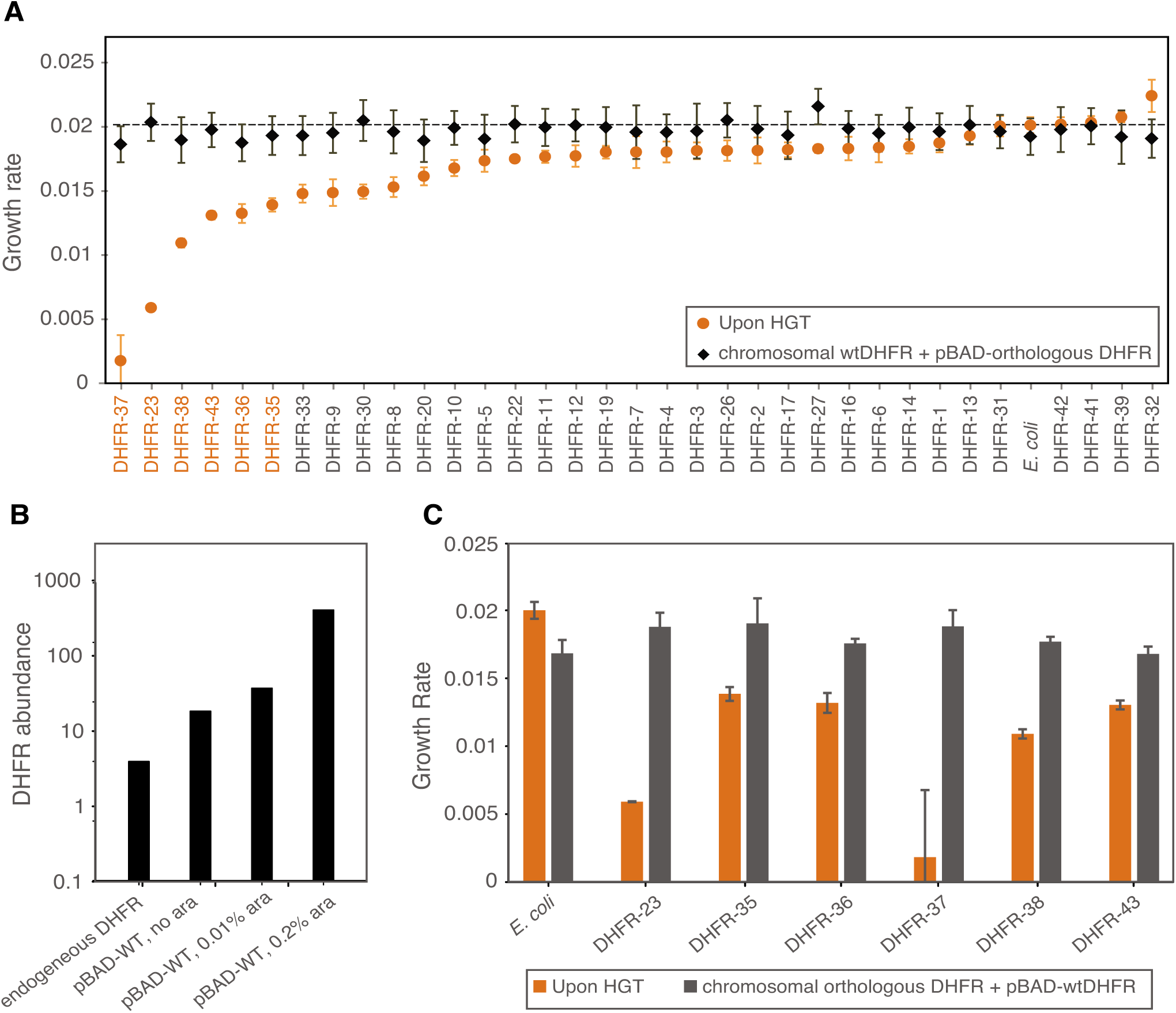
Expression of orthologous DHFR proteins is not toxic to *E. coli* cells. A) Overexpression of orthologous DHFR proteins from pBAD plasmid does not compromise fitness of WT *E. coli* strain (black diamonds). Expression was induced with 0.01 % arabinose (see **Materials and Methods**). For comparison, fitness of orthologous DHFR strains is shown (orange circles). Orthologous strains are sorted in an ascending order from less to most fit. Strains whose fitness is mostly affected by the orthologous replacements are highlighted in orange (see **Fig 1** and **S1 Table** for list of bacteria from which the orthologous DHFRs have originated). B) Intracellular DHFR abundance obtained from overexpressing WT DHFR protein from pBAD plasmid. At the highest concentration of the inducer (0.2% arabinose), intracellular DHFR levels reach ∼600 fold increase relatively to the endogenous DHFR level. Leaky expression (in the absence of the inducer) leads to 6-8 fold increase in the DHFR levels. C) Growth of orthologous DHFR strains with most severe fitness effect (DHFR-23, 35, 36, 37, 38, 43) was complemented by WT DHFR activity expressed from pBAD plasmid. No inducer was added to achieve WT intracellular DHFR abundance closer to the endogenous levels (see (B) and **Materials and Methods**). Error bars represent standard deviation of 4 independent measurements.

**S3 Fig.**
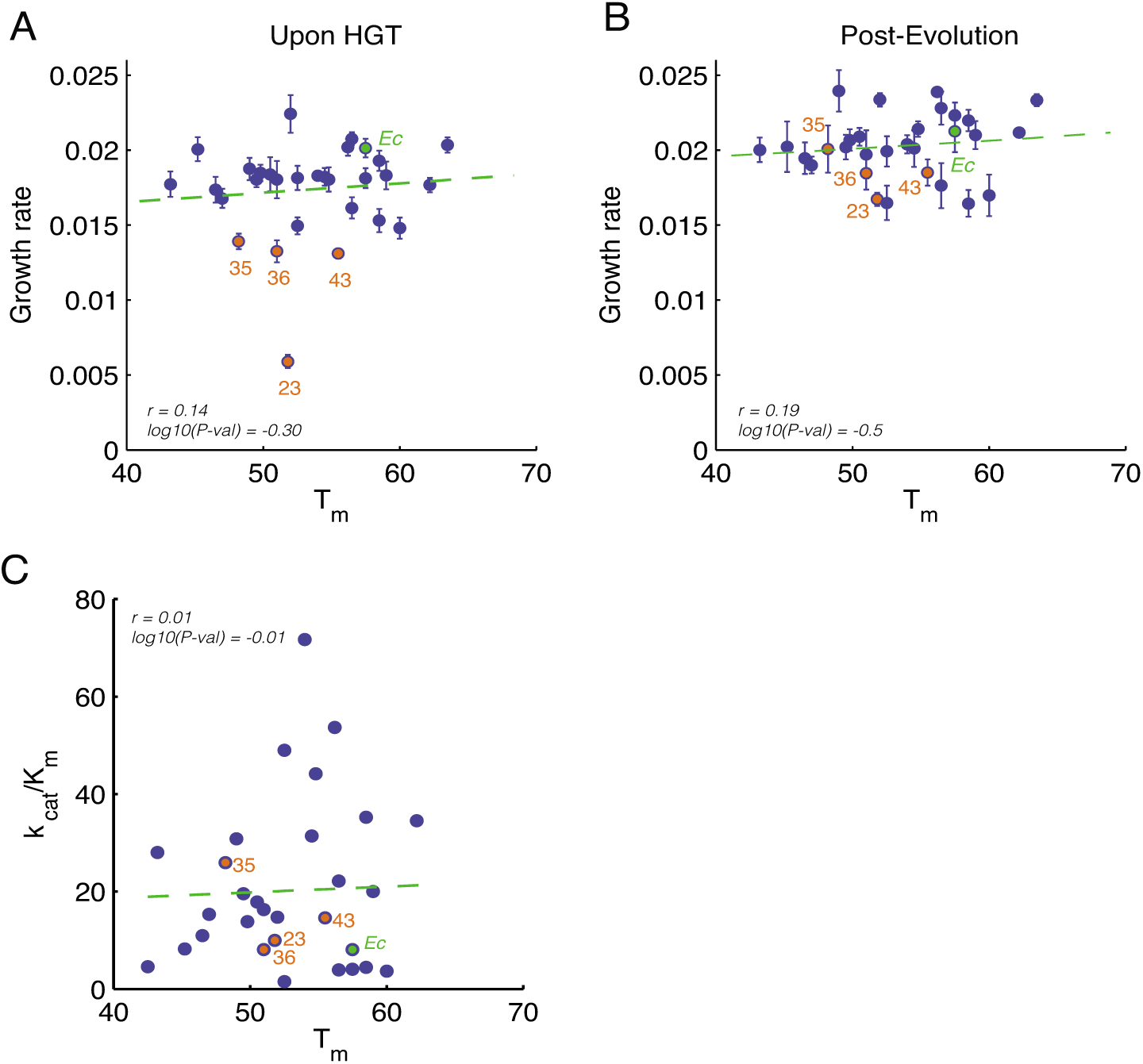
Stability of orthologous DHFR proteins does not correlate with activity or fitness. A,B) Stability of orthologous DHFR proteins (*T*_*m*_) does not correlate with growth rates of the corresponding orthologous strains before (A) or after (B) experimental evolution. C) Catalytic activity (*k*_*cat*_/*K*_*m*_) and stability (T_m_) of the orthologous DHFR do not correlate. Strains with severe fitness effects are highlighted in orange. *Ec* (green) denotes WT *E. coli* strain. Dashed green lines are regression fits to all points. Error bars represent standard deviation of 4 independent measurements.

**S4 Fig.**
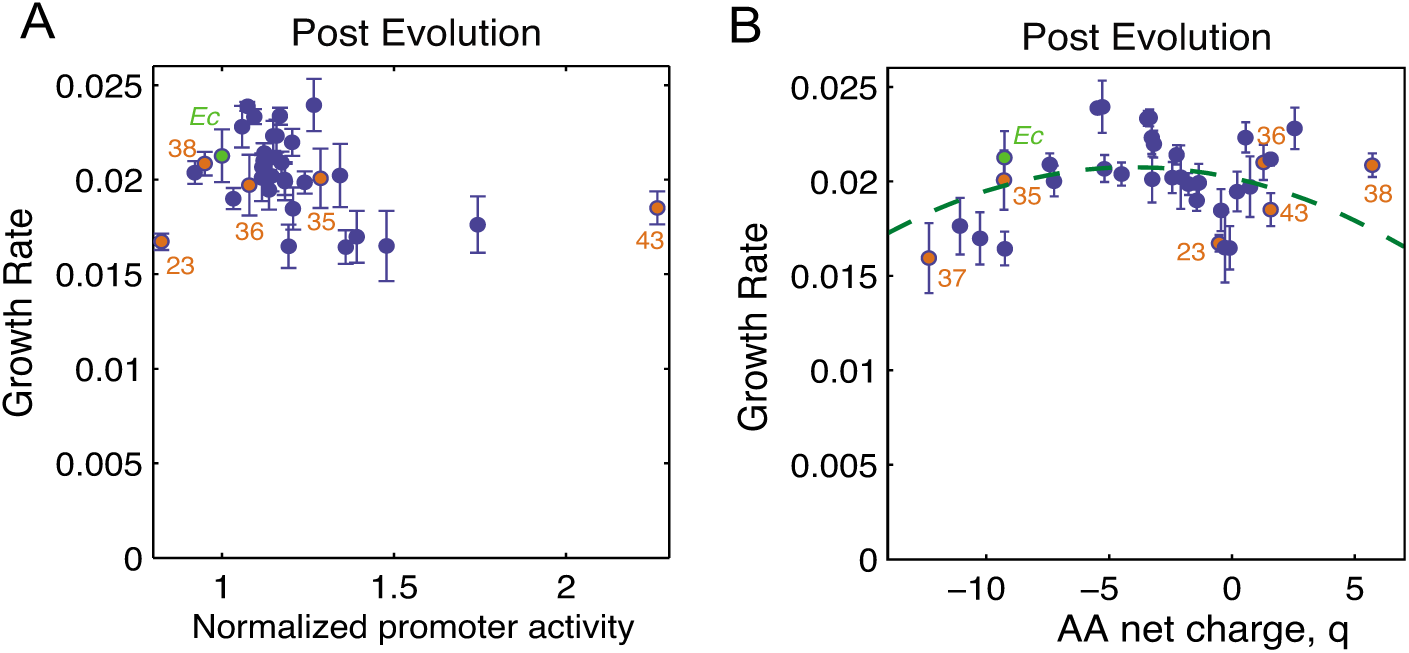
Correlation of growth rates of the evolved HGT strains with promoter activity and net charge. A) The inverse correlation between growth rate and promoter activity (**Fig 4B**) disappears after evolution. B) The significant quadratic dependence on the net charge of the DHFR amino acid sequence upon HGT (**Fig 4C**) significantly weakens after evolution. Dashed line shows the quadratic dependence to this fit (r=-0.4; p-value=0.02). Strains with severe fitness effects are highlighted in orange. Ec (green) denotes WT *E. coli* strain. Error bars represent standard deviation of 4 independent measurements.

**S5 Fig.**
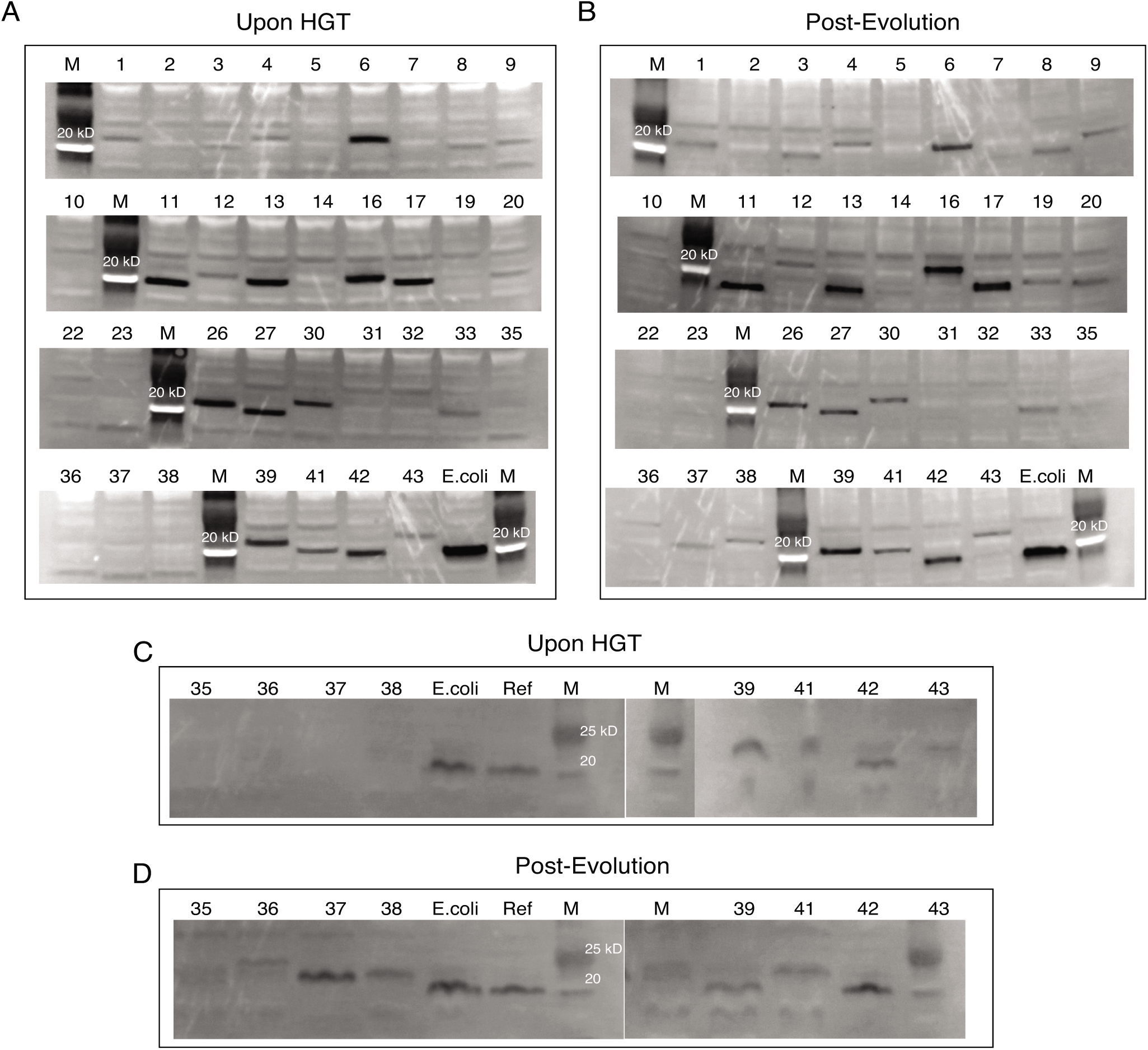
Intracellular abundance of orthologous DHFR proteins. Soluble intracellular abundances of DHFR proteins before (A) and after (B) experimental evolution. Total intracellular abundances of DHFR proteins before (C) and after (D) evolution. Numbers represent the corresponding DHFR strains (see **Fig 1** and **S1 Table**). M, marker. Ref, soluble lysate from WT strain.

**S6 Fig.**
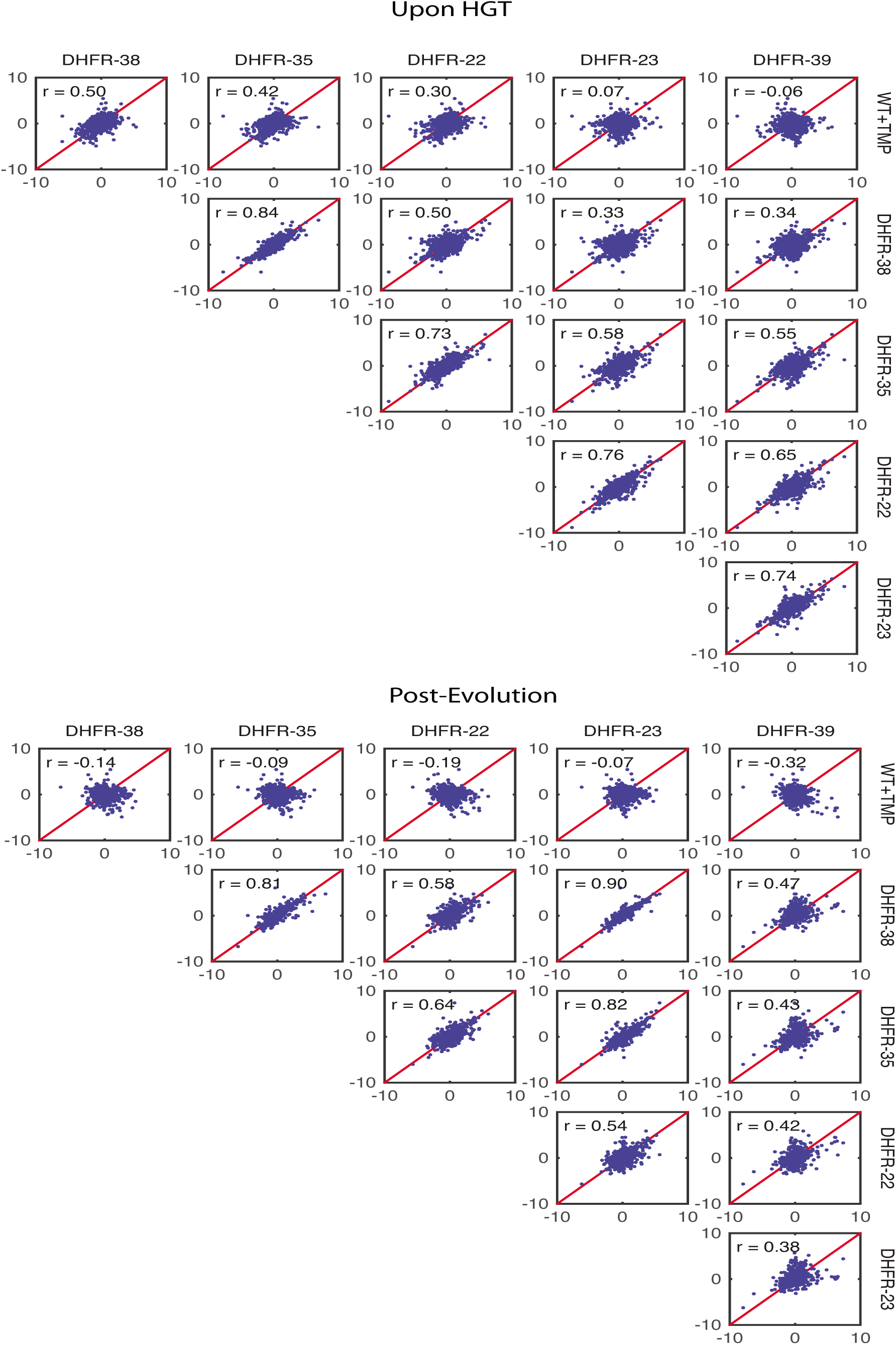
Proteomics analysis of orthologous DHFR strains. Z-score correlation plot for proteomes of orthologous DHFR-22, 23, 35, 38, and 39 strains obtained upon HGT and after evolution experiment (see **S6 Table**).

**S1 Table. Molecular properties of orthologous DHFR proteins.**

(XLSX)

**S2 Table. Codon-usage of *E. coli’*s *folA* gene and adapted DNA sequences.**

A. Aminno acids and adapted DNA sequences of orthologous DHFRs
B. Codon’s frequency of *E. coli*’s *folA* gene

(XLSX)

**S3 Table. Intracellular abundance, *folA* promoter activity and growth rates of the HGT strains.**

(XLSX)

**S4 Table. Whole-genome sequencing.**

(XLSX)

**S5 Table. Validation of whole-genome sequencing results by direct PCR.**

(XLSX)

**S6 Table. Global proteome quantification by TMT LC-MS/MS.**

A. Relative abundance
B. log10 of relative abundance
C. z-scores of log10 of relative abundance(XLSX)

